# A dynamic occupancy model for interacting species with two spatial scales

**DOI:** 10.1101/2020.12.16.423067

**Authors:** Eivind F. Kleiven, Frédéric Barraquand, Olivier Gimenez, John-André Henden, Rolf A. Ims, Eeva M. Soininen, Nigel G. Yoccoz

## Abstract

Occupancy models have been extended to account for either multiple spatial scales or species interactions in a dynamic setting. However, as interacting species (e.g., predators and prey) often operate at different spatial scales, including nested spatial structure might be especially relevant to models of interacting species. Here we bridge these two model frameworks by developing a multi-scale, two-species occupancy model. The model is dynamic, i.e. it estimates initial occupancy, colonization and extinction probabilities—including probabilities conditional to the other species’ presence. With a simulation study, we demonstrate that the model is able to estimate most parameters without marked bias under low, medium and high average occupancy probabilities, as well as low, medium and high detection probabilities, with only a small bias for some parameters in low-detection scenarios. We further evaluate the model’s ability to deal with sparse field data by applying it to a multi-scale camera trapping dataset on a mustelid-rodent predator-prey system. Most parameters are estimated with low uncertainty (i.e. narrow posterior distributions). More broadly, our model framework creates opportunities to explicitly account for the spatial structure found in many spatially nested study designs, and to study interacting species that have contrasting movement ranges with camera traps.

## 1 Introduction

Much of the data available to ecologists consists of species occurrences, which in turn have sparked the development of statistical models to analyse such data (Bailey *et al*., 2014). Due to their ability to model species occurrences while accounting for imperfect detection, occupancy models have become widely used in ecology (Bailey *et al*., 2014; Guillera-Arroita, 2017). Initial formulations of occupancy models estimated species occupancy across multiple sites that were assumed to be spatially independent (MacKenzie *et al*., 2002). However, this assumption is rarely met in the field (Johnson *et al*., 2013), and failing to account for spatial dependencies will lead to overconfidence in estimated uncertainties, and might in some cases lead to bias in estimated effects of predictor variables (Guélat & Kéry, 2018).

There are numerous extensions of occupancy models to incorporate spatial dependencies. In static occupancy models, occupancy can be made dependent on the occupancy probability of neighboring sites (Bled *et al*., 2011; Eaton *et al*., 2014; Yackulic *et al*., 2014; Broms *et al*., 2016), while in dynamic models (i.e., models that explicitly estimate change over time), colonization probability can be made a function of latent occupancy status at nearby sites. Spatial dependencies may be formulated as explicit functions of distance or connectivity between sites (Sutherland *et al*., 2014; Chandler *et al*., 2015), or in the form of random spatial effects (Johnson *et al*., 2013; Rota *et al*., 2016b).

Data from many ecological studies exhibit multiple nested spatial scales, which mirrors the fact that population dynamics result from different processes occurring at multiple scales (Baumgardt *et al*., 2019). Accordingly, recently developed multi-scale occupancy models enable analyses of data from designs with such a hierarchy of spatial scales (Nichols *et al*., 2008; Aing *et al*., 2011; Mordecai *et al*., 2011; Kéry & Royle, 2015; Smith & Goldberg, 2020), and can be extended to dynamic versions to estimate colonization and extinction probabilities (Tingley *et al*., 2018).

A parallel development of occupancy models—dynamic multi-species models— addresses how interacting species co-occur over time (MacKenzie *et al*., 2004; Waddle *et al*., 2010; Richmond *et al*., 2010; Rota *et al*., 2016a; MacKenzie *et al*., 2017; Fidino *et al*., 2019; Marescot *et al*., 2020). These models have great potential to increase our knowledge of species interactions. However, interacting species in general, and predators and prey in particular, often move at different spatial scales (de Roos *et al*., 1998; Fauchald *et al*., 2000). Incorporating the multiple spatial scales of interacting species would therefore lead to more intuitive and ecologically meaningful model parameters as colonization and extinction parameters may represent different ecological processes on different spatial scales.

Here we build on dynamic multi-species models by MacKenzie *et al*. (2017) and Fidino *et al*. (2019), as well as the dynamic multi-scale occupancy model by Tingley *et al*. (2018) to develop a multi-scale dynamic two-species occupancy model. In this model, initial occupancy, colonization and extinction probabilities are estimated at two spatial scales, i.e. both at a site level and at a block level, spanning a cluster of sites. After describing the model, we perform a simulation study to investigate potential issues of bias and precision under different scenarios. Finally, we apply the model to a camera trapping data set with two spatial scales to estimate the predator-prey interaction strength between small mustelids and small rodents.

## 2 Model

### 2.1 Motivating example

The case study that motivated the development of this model was a camera trap dataset from the long-term monitoring program COAT (Climate-ecological Observatory for Arctic Tundra, Ims *et al*., 2013). The camera trapping program targets small rodents and their small mustelid predators. Small rodents (grey-sided vole *Myodes rufocanus*, tundra vole *Microtus oeconomus* and Norwegian lemming *Lemmus lemmus*) constitute prey, while small mustelids (stoat *Mustela erminea* and least weasel Mustela *nivalis*) constitute predators. Rodents and mustelids have for decades been known to exhibit a predator-prey interaction (Hanski *et al*., 1991), which is one of the main hypotheses for the population cycles of rodents (Krebs, 2013). However, estimating the prevalence and strength of that interaction has proved difficult, as reliable data—especially on mustelids—have previously been lacking (King & Powell, 2006). Recently camera traps have been tailored to monitor small mammals, including rodents and mustelids (Soininen *et al*., 2015). These camera traps have been demonstrated to be functional year-round in Arctic tundra habitats (Mölle *et al*., 2021), whereas data was previously lacking in winter. The sampling design of the monitoring program has a multi-scale structure, where sites (camera traps) are spaced >300m apart, but clustered in blocks of 11 to 12 cameras covering two different habitats, snowbeds and hummock tundra (see Appendix S1, section 3). This spatial structure can be matched to the movement ranges of small rodents and mustelids (Hellstedt & Henttonen, 2006), where sites represent independent samples of rodent presence and blocks represent independent samples of mustelid presence. The camera trap monitoring was started in the autumn of 2015 and consisted of 4 blocks, with 11 camera trap sites within each block. In the summer of 2018, the monitoring was expanded by 4 blocks, containing 12 camera trap sites each, to make up a total of 8 blocks. We here aim to investigate the strength of the interaction between rodents and mustelids using this camera trapping dataset. The spatial hierarchy in the sampling design leads to spatial dependencies in the data that needs to be accounted for. However, it also provides an opportunity to investigate the predator-prey interaction on two nested spatial scales. The case study is viewed as an inspiration for the following model framework, which is nonetheless more general.

### 2.2 Model description

#### 2.2.1 Model structure and latent ecological states

Our dynamic occupancy model for interacting species has two spatial levels (see Fig. 1 for an illustration of the spatial design and the model structure). On the block level the model has *B* blocks (*b* ∈ {1,…,*B*} being the index of the blocks), each containing *K* sampling sites (*k* ∈ {1,…,*K*} being the index of the sampling sites). Each sampling site is surveyed over *T* (primary) sampling occasions, *t* ∈ {1,…, *T*} being the index of the primary occasion. Between the primary occasions the populations are assumed to be open (i.e. the species can colonize or go extinct). Within each primary occasion each site is sampled *J* (secondary) occasions, *j* ∈ {1,…, *J*} being the index of the secondary sampling occasions. Between the secondary occasions the populations are assumed to be closed (the species are assumed to neither colonize nor go extinct at any site). For each secondary occasion j, considering two species (or functional group), any given site can then either be observed as unoccupied (*y_b,k,t,j_ = U*), occupied by species A only (*y_b,k,t,j_ =* A), occupied by species B only (*y_b,k,t,j_ = B*) or occupied by both species A and B (*y_b,k,t,j_ = AB*). The replicated samples within primary occasions, during which the populations are assumed to be closed, allow for the estimation of the detection process (MacKenzie *et al*., 2017). The true latent state of each site (*k*) during each primary occasion (*t*) in a given block (*b*) can then be described with a latent variable *z_b,k,t_*, that can take on any of the same 4 states. In this model we also consider an ecological process on the block level by describing a latent block state, *x_b,t_*, that can take on any of the same four states as the latent site level state (U, A, B, or AB). This process occurs on a larger spatial scale than the site level, and will for instance in our case study represent the spatial scale of dispersal (e.g. changing home ranges) for mustelids, compared to the site level which represents the foraging movements within their home range. The site states will then depend on the state of the block they belong to (e.g. if a block is unoccupied by a given species, all sites within that block also have to be unoccupied by that species, although the converse is not true).

**Figure 1:**
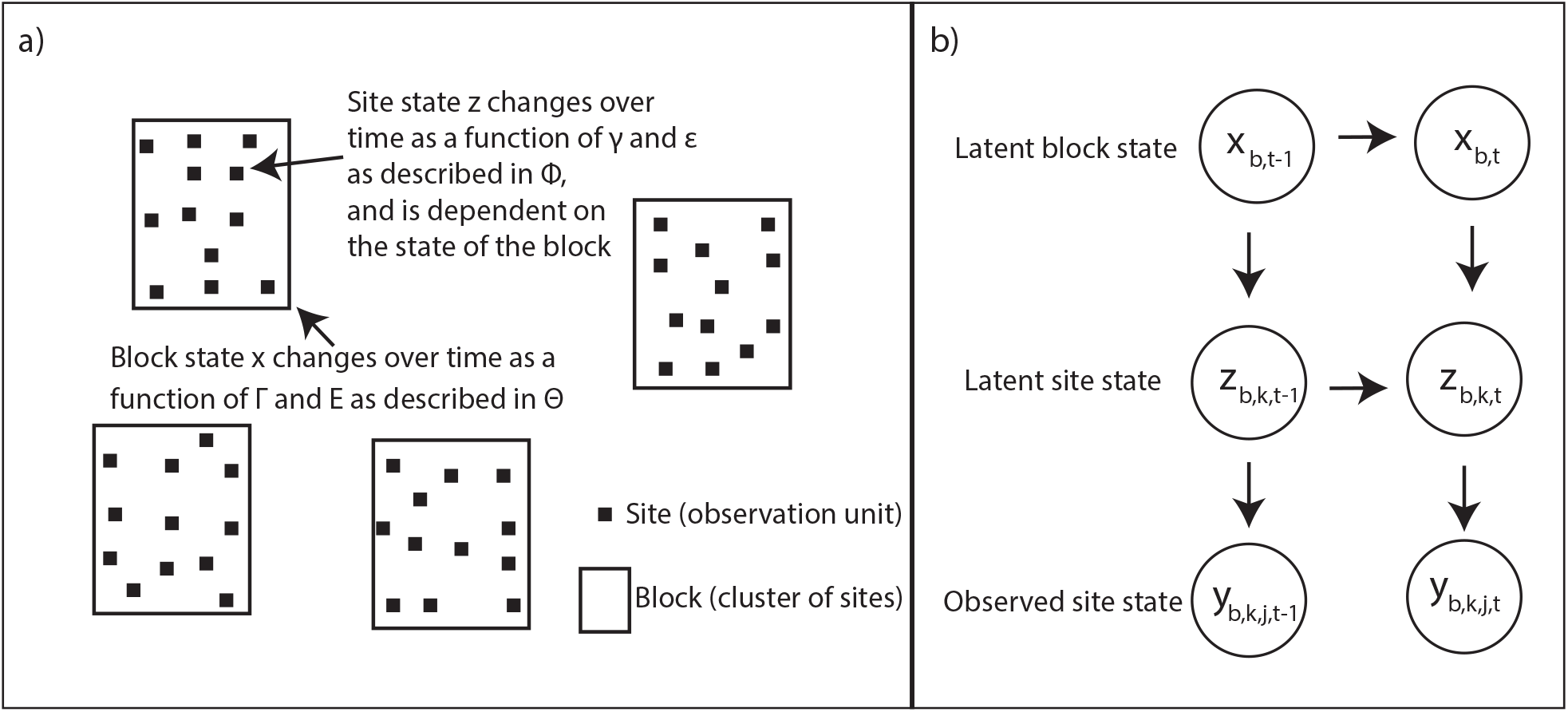
Conceptual diagram of the design and model structure. Panel (a) describes the multi-scale structure of the design, data sampling and state variables (probabilities) with clusters of sites nested in 4 blocks. Parameters *γ* and *ε* denote site level colonization and extinction probabilities, respectively, and Φ denotes the site transition probability matrix. Parameters Γ and *E* are the block level colonization and extinction probabilities, respectively, and Θ denotes the block level transition probability matrix. The diagram in panel (b) shows the conditional dependencies among the state variables in the model for observation j in primary occasion t at site *k* in block b.

#### 2.2.2 Transition model

After initial states (i.e. the site and block level states in the first primary occasion) have been modelled as a random categorical variable (see Appendix S1, section 1 for details), transitions between states are modelled with the transition probability matrices Θ for the block level and Φ(***x**_b,t_*) for the site level, with the latter depending on the block state (see Fig. 1 for an illustration of the spatial setup and the model structure).

It is assumed that the site colonization (*γ*) and extinction (*ε*) probabilities are dependent on the site state in the previous time step (*z*_*k,b,t*–1_) as well as the block state in the same time step (*x_b,t_*). The latter is assumed because site level colonization is only possible whenever the given species is already present in the block. Because site transition probabilities are dependent on the block state in the same time step (Φ(*x_b,t_*)), it is possible that both a block and sites within that block are colonized in the same time step. However, because sites do not cover all available habitat within the block, blocks are not forced to go extinct even though all sites within that block are extinct. Hence, the model allows for a block to remain occupied even when all of its sites go extinct. On the other hand, when a block goes extinct, all of its sites are forced to go extinct (i.e. if *x_b,t_* = U then *z_b,k,t_* = U for all *k* ∈ {1,…,*K*}). Overall, the block-level latent states follow a classical Markov chain where the latent states at *t* are only determined by the latent states at *t* – 1 while the site latent states follow a Markov chain where the latent states at *t* depend on the site latent states at *t* – 1 and the block state at *t*: there is a forcing of the site-level state transitions by the current block state but otherwise the latent states follow a Markov chain between primary occasions. For that reason, it is important that the temporal resolution of the primary occasions correspond to the time scale relevant for the interaction of interest.

The transition probability matrix (Θ) for blocks (*b*) can be written as follows, with verbal definitions of block level transition parameters given in Table 1,

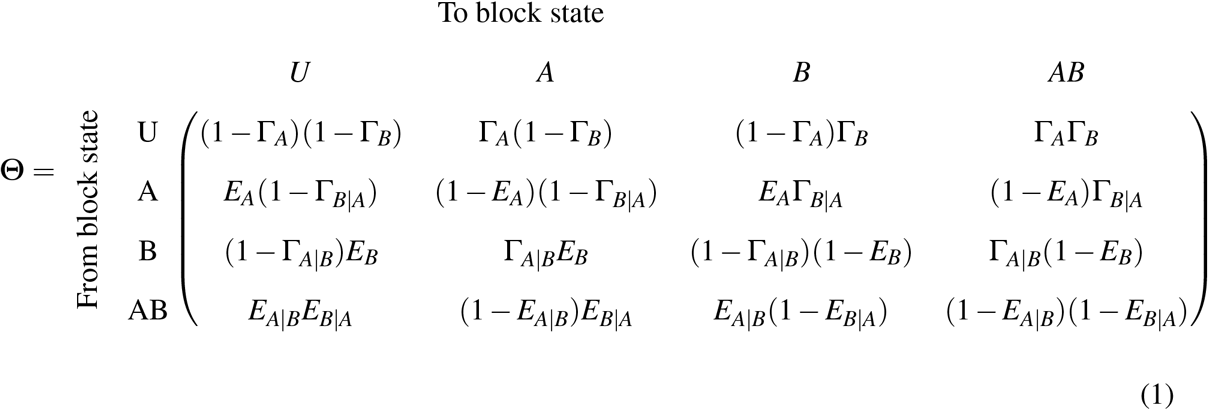

**Table 1:**
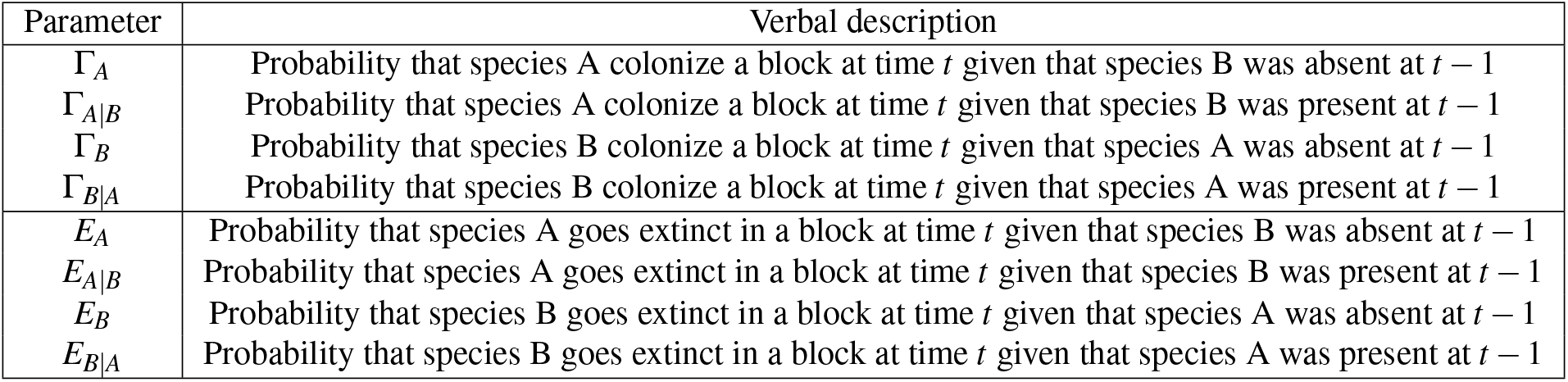
Verbal definition of block level transition parameters.

Since the site transition probabilities depend on the block level state (*x_b,t_*) we create one site transition matrix for each possible block state in the following way with verbal definitions of site level transition parameters given in Table 2:

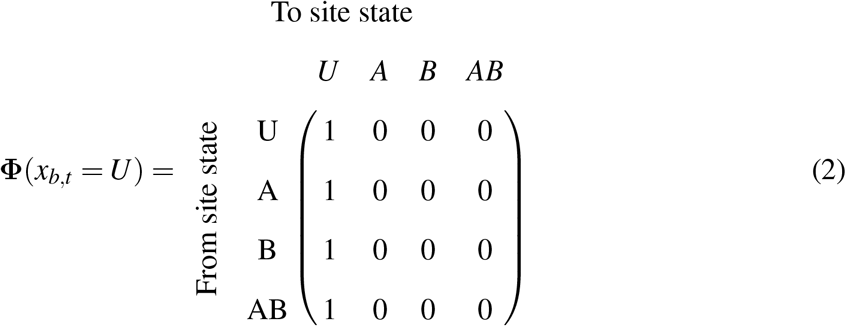

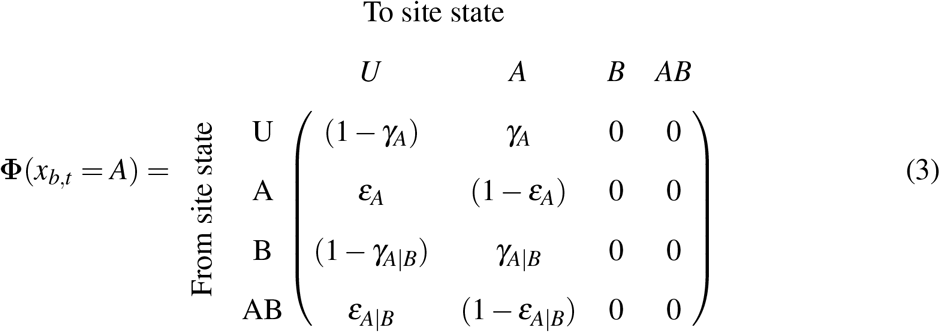

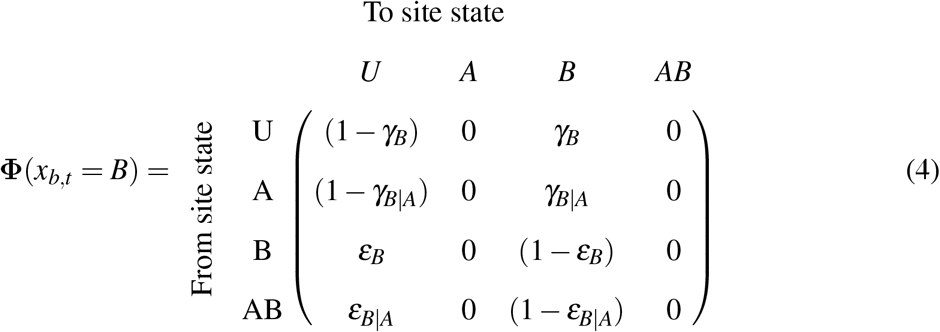

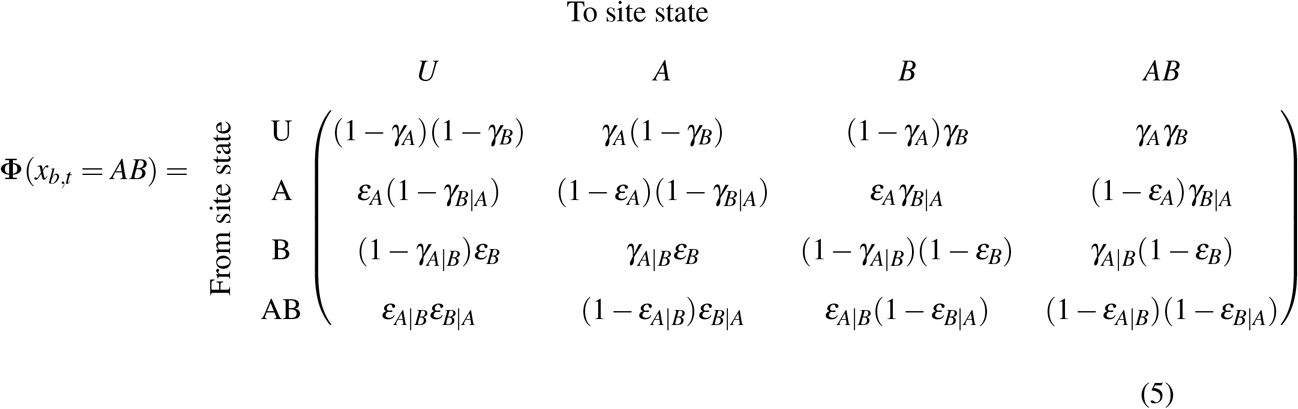

**Table 2:**
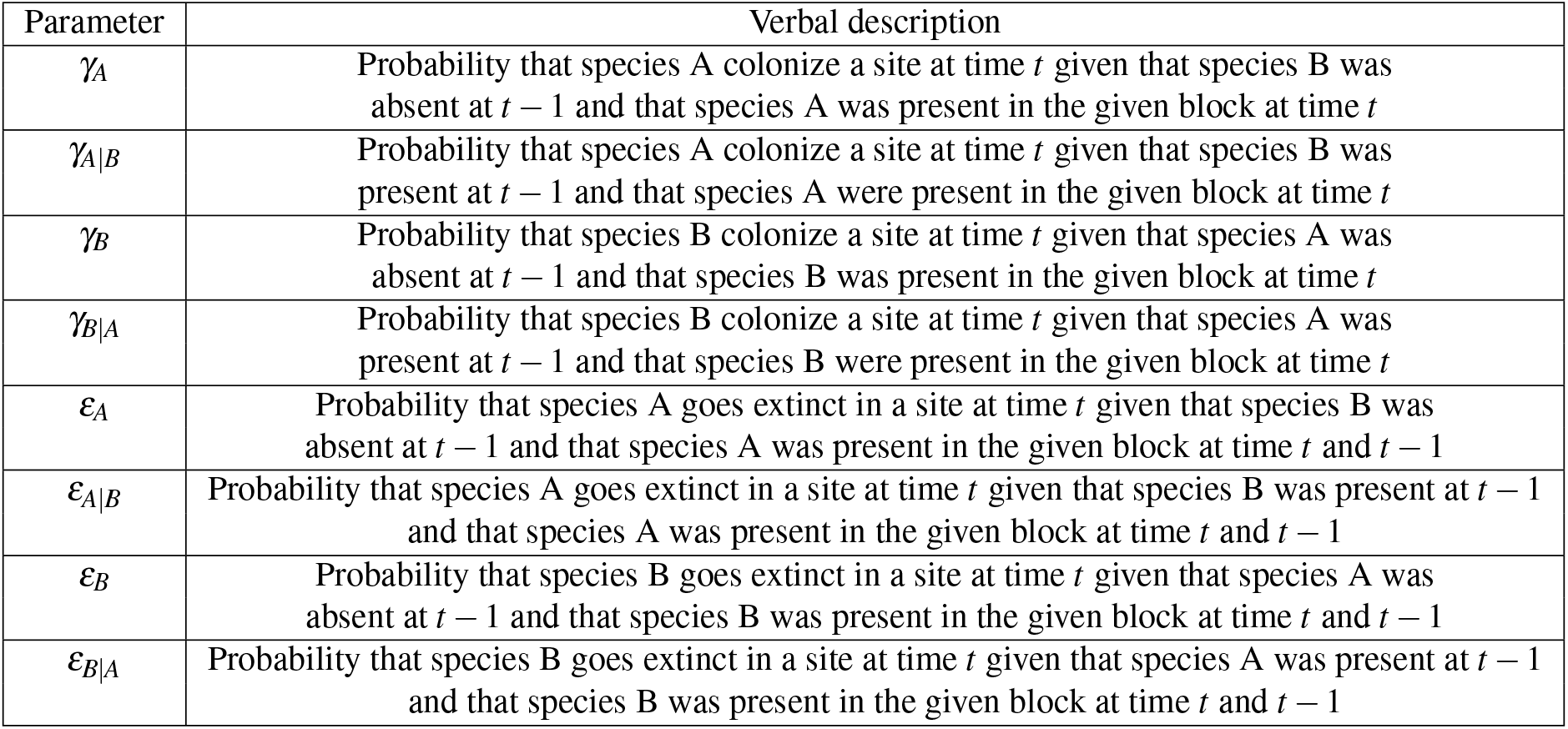
Verbal definition of site level transition parameters.

Then the full model for both block- and site-level states can be written as

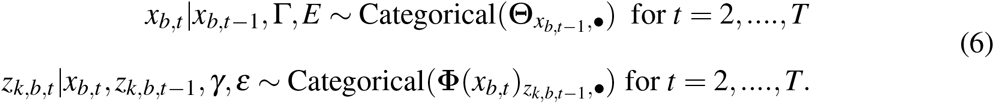

The indices *x*_*b,t*–1_ describe the specific row in Θ which has value *x*_*b,t*–1_ and *z*_*k,b,t*–1_ the row in Φ which has value *z*_*k,b,t*–1_, while ● refers to all columns of that row.

#### 2.2.3 Detection model

All observations on which the model relies on are coming from the site level. Hence, the detection probability can be modelled similarly to other two-species occupancy models. Models to estimate detection probabilities of one species dependent on the presence (Rota *et al*., 2016a) or the detection of the other species (Miller *et al*., 2012; Fidino *et al*., 2019) exist. It is possible that such detection-interactions also exist for mustelids and rodents. However, as we have no indications that they do, for simplicity, we assumed that the detection of each species is independent of both the presence and detection of the other species. In the simulation study we also assumed that the detection probability is constant over sites, blocks and temporal occasions. However, note that a temporal binary covariate is added to the detection model in the empirical case study (see related section below). Let *p_A_* and *p_B_* be the detection probabilities of species A and B at a given site, the detection probability matrix (*λ*) can then be defined as

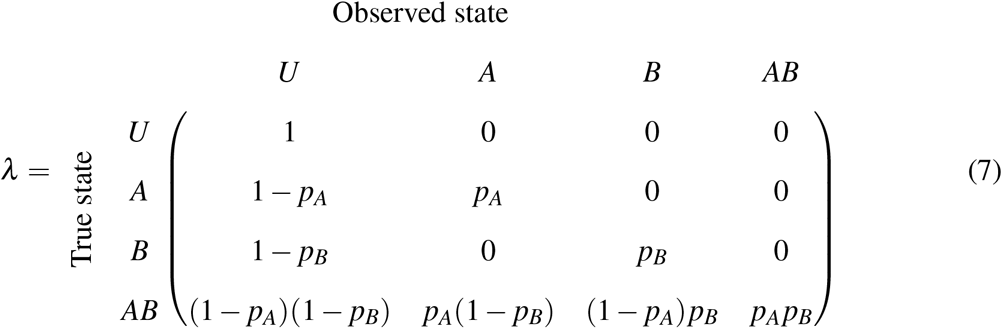

The observation at site *k* in block *b* at visit *j* during time *t* (*y*_*k,b,j,t*_) can then be described by the following equation:

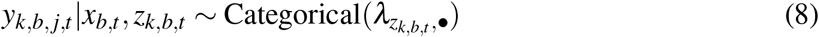

with *z_k,b,t_* being the chosen row of the detection matrix (*λ*) to draw the observed state.

#### 2.2.4 Model fitting

The models were analyzed in a Bayesian framework using the JAGS software (Plummer, 2003) with the `jagsUI` package (Kellner, 2015) in `R v4.0.3` (R Core Team, 2020). We note that the model is a hidden Markov model and could also be fitted in a maximum likelihood framework using the Forward algorithm (McClintock *et al*., 2020). Convergence was assessed by having a 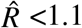 for all key parameters (Gelman *et al*., 2013) and from graphical investigation of traceplots.

## 3 Simulation study

We conducted a simulation study to evaluate the performance of our model by examining potential issues of bias and variance in parameter estimates of colonization, extinction and detection. The spatiotemporal structure of the simulated data was largely inspired by our empirical case study (see section below for a more detailed explanation). Hence, we simulated data for 8 blocks, each containing 12 sites. We chose weeks as primary occasions and days within each week as secondary occasions corresponding to the expected rate of the dynamics of the empirical case study (An-dreassen & Ims, 2001). We simulated data for 50 weeks (approximately one year). The chosen parameters values were also inspired by the predator-prey case study (e.g. *E_A_* <*E_B_*; see all parameter values in Appendix S1). To investigate how contrasting scenarios affected bias of parameters, we simulated data where both species had low (lo), medium (mo) and high (ho) average occupancy probability (with medium detection probability). In addition we simulated data where both species had low (ld), medium (md) and high (hd) detection probabilities (with medium occupancy probability, for more details, see Appendix S1, section 2). For each scenario, we simulated and analyzed 50 replicate datasets. We specified Uniform(0,1) prior distributions for the detection, colonization and extinction probabilities and Uniform(0,0.5) prior distributions for the initial occupancy probabilities. The MCMC algorithm was run with an adaptation phase (initial phase where the Bayesian sampler can adapt to increase efficiency) of 1000 iterations. No additional iterations was discarded as burn-in. The model was run for 5000 iterations with a thinning of 10 (keep every 10 values in the chain to construct the posteriors).

### 3.1 Results from the simulation study

All colonization and extinction probabilities at both spatial scales were estimated without bias for most data scenarios (Fig. 2 and Tables S5-S7 in Appendix S1). However, in the low detection scenario there was a slight positive bias in the block extinction probability of species B when species A was absent (*E_B_*, 21% bias) and in the block extinction of species A when B was present (*E*_*A*|*B*_, 20% bias, see Fig. 2). Both detection probabilities (*p_A_* and *p_B_*) were estimated without any apparent bias (Fig. S11 in Appendix S1). The initial values were also estimated without any obvious bias except for *ψ_AB_* under the low occupancy and low detection scenario (Fig. S12 in Appendix S1).

**Figure 2:**
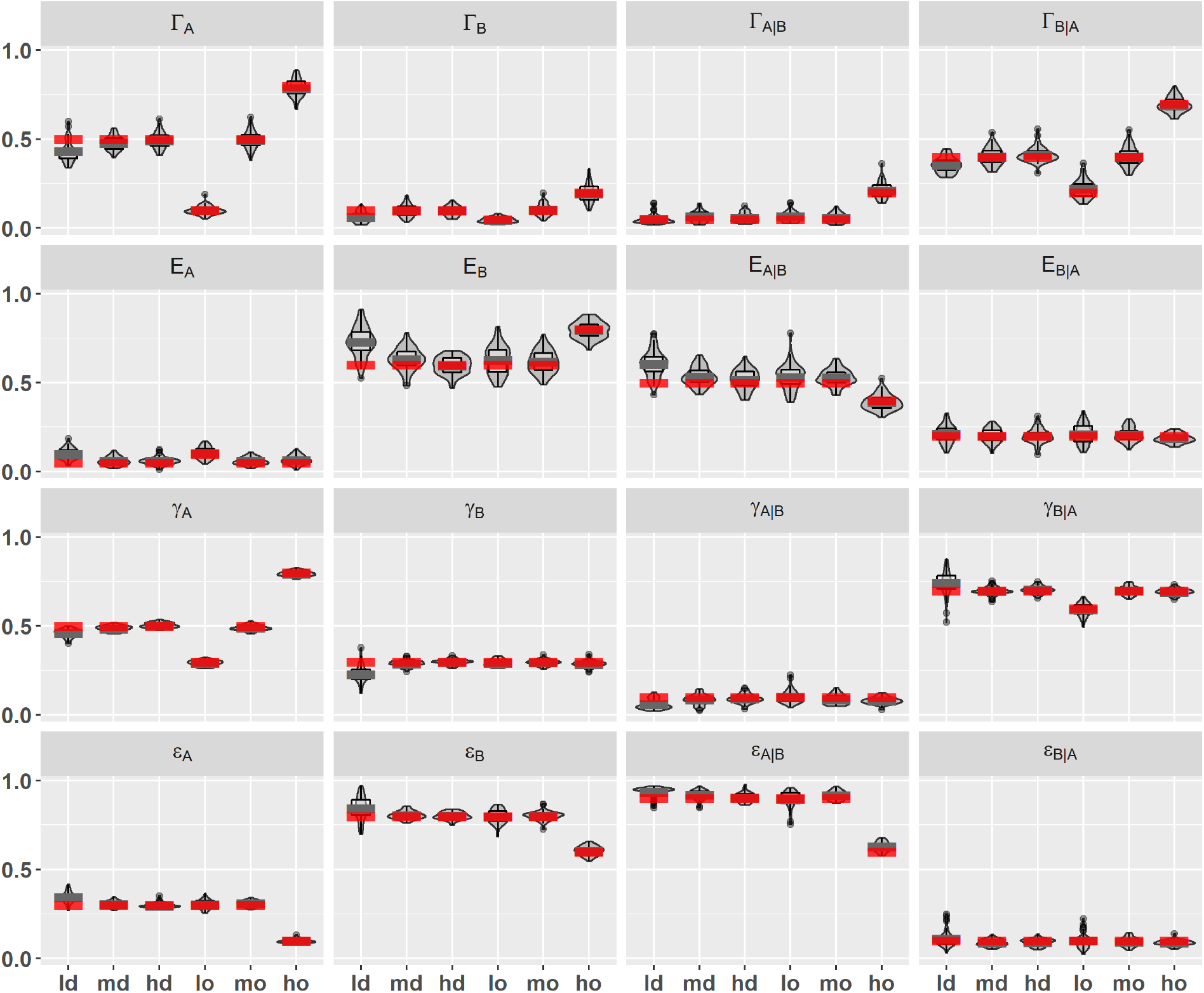
Violin plots and boxplots of the posterior means of site and block colonization (*γ* and Γ) as well as extinction probabilities (ε and *E*), from 50 simulation replicates. The thick red bar indicates the true parameter values while the thick grey bar indicates the average of the posterior means from the 50 simulated replicates. The x-axis displays the 6 different data scenarios: low, medium and high detection probability of both species (ld, md, hd) and low, medium and high average site occupancy probability of both species (lo, mo, ho).

## 4 Empirical case study

Working with camera trap data requires to associate species identity to the photographs taken by cameras. For this we used the automatic classification model of Tabak *et al*. (2019) implemented in the MLWIC R-package (Tabak *et al*., 2018, see Appendix S1 section 3 for more details). As there were particularly few observations of least weasel and Norwegian lemming, the analysis could not be conducted at the species level. However, as both mustelid species prey on all rodent species present in tundra, rodents and mustelids were considered functional groups. As the rodents are known to exhibit rapid local scale colonization-extinction dynamics (Andreassen & Ims, 2001) we here define a primary occasion as one week (i.e. 7 days) and secondary occasions as the days within that week. While mustelids have slower demographic processes than small rodents, they are assumed to show a spatially aggregative response (movement towards prey-rich areas) on a short time scale (i.e., changing foraging grounds from one week to the next, Hellstedt & Henttonen, 2006). The resulting dataset describes, for each day, if each functional group was detected or not. We combined the data for the two functional groups in a multi-state occupancy dataset with 4 states (U = none of the species are observed, A = only small rodents observed, B = only small mustelids observed or AB = both small rodents and small mustelids are observed). We included all weeks from the beginning of the monitoring in 2015 to the summer of 2021, resulting in a total of 304 weeks. Note that the different blocks were established at different times. Hence, since more than half of the sites were only observed for the last 47 weeks this resulted in the inclusion of some missing data in the earlier years. Since mustelid observations mostly occurred in the snowbed habitat, we chose to focus only on these sites, reducing the number of sites to 5 - 6 per block.

This study system exhibits strong seasonality that needs to be accounted for. However, there is a lack of detailed environmental data representative of the seasonality in this area. Therefore, we used temperature measurements from the camera traps to estimate the onset of winter (defined as the first day after summer with a daily mean temperature < 0) and the time of snow melt (defined as the first day in spring with daily mean > 0) for each camera trap individually. Such partitioning of the year will account for most of the seasonal changes affecting detection, as the winter period is mostly snow-covered, forcing small mammals into the subnivean space at the bottom of the snow pack, while the summer season is mostly snow-free. Indeed, presence of snow is known to impact the detection probability of small rodents on the Arctic tundra (Mölle *et al*., 2021). Season was therefore included as a covariate on detection probabilities (*p_A_* and *p_B_*) through a logit-link function. The model was analyzed similarly to the simulation model.

We performed a prior sensitivity analysis by running the model with 3 different sets of priors. The first set contained flat uniform priors (~ Uniform(0,1)) except for the detection probabilities, whose intercepts and slopes were given Normal(0,1) prior distributions since they are defined on a logit scale. For the second set we used centered priors by using a Beta(4,4) distribution for all priors, again except for the detection probabilities, which now were given a Logistic(0,1) prior distribution. The third set of priors was specified to be skewed towards our ecological expectations, by using either a Beta(2,4) or Beta(4,2) depending on the expected relationships in a predatorprey system. Detection probabilities were given a Normal(−0.5,1) prior distribution on the logit scale (see Appendix S1, section 4). The prior sensitivity analysis was also used to investigate identifiability by additionally estimating prior-posterior overlap (Gimenez *et al*., 2004).

To assess the goodness of fit we calculated a Bayesian p-value (*p*_Bayes_) as described by Kéry & Royle (2020) (see details in Appendix S1, section 4) and performed graphical predictive checks. To reach convergence with the field dataset we needed to run the model for 20 000 iterations, with 5 000 steps of adaptation. In addition we discarded the first 1000 iterations as burn-in and used a thinning of 20. We note that this was slightly longer than for the simulation study.

### 4.1 Model performance

The posterior distributions for all parameters are similar for the 3 sets of priors. The estimated posterior distributions are also consistently different from the prior distributions. Therefore, it appears that all colonization and extinction parameters on both spatial levels are identifiable from the data. Regarding goodness-of-fit tests, we obtained more mixed results. For the open (i.e. dynamic) part of the model, there is a reasonable fit to the data for both functional groups (see Appendix S4 for graphical predictive checks—although Bayesian p-values are not very close to 0.5, *p*_Bayes_ for rodents = 0.37 and *p*_Bayes_ for mustelids = 0.91). For the closed part of the model (i.e. detection process), the model fit seems to be poor for mustelids and even more so for rodents. This could indicate that there are factors affecting the detection probabilities that we do not account for, or that the discretization of the continuously sampled camera trap data induces some lack of fit. We stress, however, that models checks could be further improved upon. For instance, the Bayesian p-values for the closed part of the model calculated on the simulated data (where model fit is known a priori to be correct) were actually already close to 1 rather than 0.5, which means that they are not adequately doing their job. We relied instead on graphical comparisons of predicted vs observed scores, which also suggested a lack of fit for the closed part of the model.

### 4.2 Results from the empirical case study

According to the fitted model rodents generally had a higher detection probability than mustelids, and both species were slightly less detectable during winter than in summer (for rodents: 0.53 with 95% CRI (0.51-0.54) during winter and 0.59 with 95% CRI (0.58-0.60) during summer, for mustelids: 0.14 with 95% CRI (0.11-0.17) during winter and 0.16 with 95% CRI (0.14-0.18) during summer).

The strongest evidence for predator prey-interactions was found in the estimated extinction probabilities on the site level (Fig. 3), where mustelid presence increased the rodent extinction probability from 0.13 to 0.34 (an increase of 0.21 with 95% CRI (0.13 - 0.29)). More surprisingly, mustelid extinction probability on the site level also increased (0.22 with 95% CRI (0.11 - 0.35)) in the presence of rodents. There was no site-level evidence for effects of predator-prey interactions on the colonization probabilities, nor was there evidence for effects of predator-prey interactions on any of the block level parameters (Fig. 3). However, it must be noted that the estimated probabilities at the block level were small and/or uncertain (Table S9 in Appendix S1). Especially, we note that block-level colonization of rodents when mustelids were present (Γ_*A*|*B*_) and block-level extinction of mustelids alone (*E_B_*) were estimated with large uncertainties (Fig. 3). See Tables S8 and S9 in Appendix S1 for numerical values of estimates of all colonization and extinction probabilities, and the differences between them, with corresponding credible intervals.

**Figure 3:**
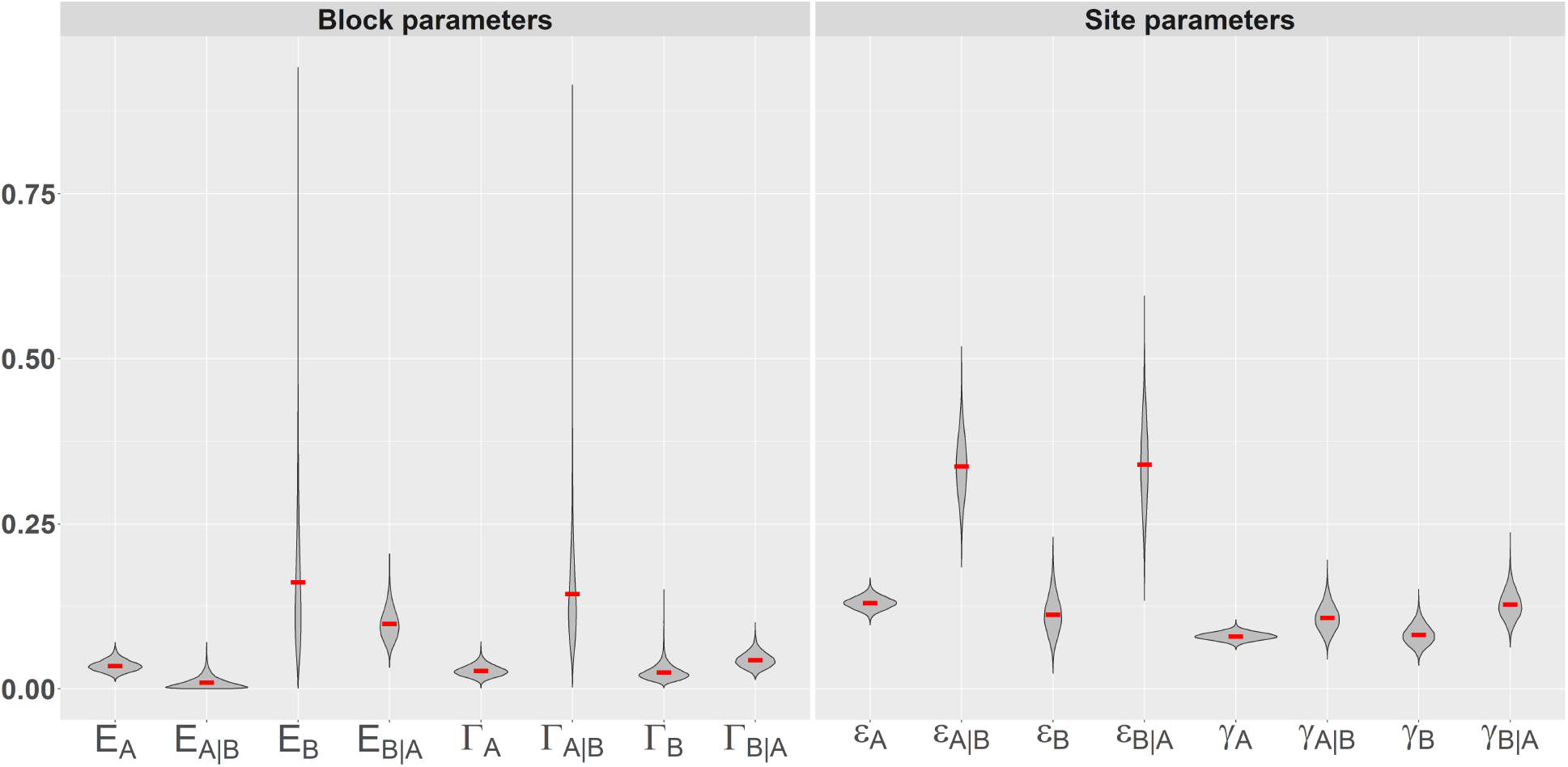
Violin plots of the estimated posterior distribution of site and block colonization (*γ* and Γ) and extinction probabilities (*ε* and *E*) for the case study. Subscript *A* denotes rodents and *B* denote mustelids. Red bars indicate posterior means.

## 5 Discussion

We constructed a dynamic multi-scale occupancy model for interacting species. Through simulations, we demonstrated that this model is able to produce unbiased estimates of colonization and extinction parameters under most scenarios of average presence and detection.

We find the current extension of the dynamic multi-species occupancy framework to nested spatial scales useful for two reasons. First, it make it possible to explicitly account for the spatial structures found in many spatially nested study designs. This should reduce bias and increase precision in parameter estimates. Second, it makes it possible to investigate the joint colonization and extinction dynamics of species pairs that have contrasting movement ranges. In case of predatorprey, the daily movement range of one species (e.g., the predator) may be equivalent to the dispersal distance for the other species (e.g., the prey). Indeed, analysing data from a multi-scale monitoring program of two species with such contrasting movements ranges and spatially nested design (as in our mustelid-rodent case study) with a single-scale model would make little sense. Not only would this violate the assumption of spatial independence of sites, but it would also compromise the ecological interpretation of parameters, that would then mix different kinds of movements (e.g. foraging and dispersal movements for the predator). By contrast, our dynamic multi-scale model is able to disentangle the scale-dependent nature of colonization and extinction probabilities, in our case, at the site and the block level. Our work therefore helps to bridge the gap between the separate developments of multi-scale dynamic occupancy models (Tingley *et al*., 2018) and dynamic multi-species occupancy models (MacKenzie *et al*., 2017; Fidino *et al*., 2019).

The simulation study showed that all model parameters are estimated without any considerable bias in most data scenarios (Fig. 2, see Appendix S1 Table S6 for numeric values). The exception is the low detection scenario where small biases appeared for the block extinction probability of A when B is present (*E*_*A*|*B*_) and the block extinction probability of B when A is absent (*E_B_*). A likely explanation is that when few blocks are occupied with species B, there are also few blocks that have the potential to go extinct. Furthermore, the block-level parameters appear to vary more between models than the site-level parameters. This could result from all observations coming from the site-level, with the block-level parameters thus depending on the reconstruction of two latent states, with a propagating uncertainty. Moreover, the nested study design has by construction more sites than blocks, providing more data to estimate the site-level parameters than the block-level parameters. If an auxiliary data stream constituted of indices of block presence (e.g. snow track data for mustelids) could be constructed, it could be fed to a joint model to increase the precision on estimated block-level parameters. In the current model framework, the simulation study demonstrates that care has to be taken when analysing data on species with low detection probability. We encourage further work on the data requirements of similarly complex multi-scale occupancy models, possibly with more informed priors and/or additional data streams. We consider this simulation exercise as opening doors rather than providing definitive answers.

Our real-world case study incorporates some of the classical empirical challenges that ecologists have to deal with. First, the dataset has a high proportion of missing data and detection probability differs between the functional groups. Second, the dataset comes from an ecosystem with both strong seasonal and inter-annual (i.e. cyclic) variability, which likely affects the detection, colonization and extinction probabilities. Third, we use functional groups instead of species, potentially adding some unexplained variability in the data. Moreover, we treated blocks as if they were identical, which is likely to be an oversimplification. Even without addressing these challenges specifically in the model (with the exception of a seasonal covariate on detection probability), the model seems to be able to identify most parameters. However, it is evident that some parameters (Γ_*A*|*B*_ and *E_B_*) are estimated with large uncertainties, to a degree where it limits the ecological inferences that can be made (Fig. 3). Why Γ_*A*|*B*_ and *E_B_* are the parameters that are estimated with largest uncertainty can be explained. First, these are block-level parameters, and by design fewer transitions between states occur at block level (since there are less blocks than sites). These are also predator-related parameters, conditional on predator presence, and there were fewer observations of mustelids than of rodents (see Table S4 in Appendix S1). It is expected that predators have fewer observations than prey as they have lower population density. However, mustelids are also known to be especially difficult to observe (King & Powell, 2006). Our empirical case study appears thus to constitute the minimal data requirements for this model. It is likely that a longer time series would increase parameter precision (Guillera-Arroita *et al*., 2014), especially in the case study system where the population dynamics are ruled by multi-year population cycles. Other predator-prey systems where the predator is more conspicuous might also make it easier to identify all parameters including at block level.

Despite the abovementioned challenges, the model gave some evidence of a predator-prey interaction between mustelids and rodents (as hypothetized in Hanski *et al*., 1993; Norrdahl & Korpimäki, 2000). On the site level, mustelid presence lead to an almost four-fold increase in rodent extinction probability, which is probably due to direct effects (killing) and indirect effects (predator avoidance) of predation by mustelids. This result is coherent with the hypotheses that mustelids are able to induce population crashes in boreal and arctic rodent populations (Hanski *et al*., 1993). Most of the current observational evidence for this hypothesis comes from indirect observations of mustelids (snow tracks Korpela *et al*., 2014 and winter nests Gilg *et al*., 2003) and spatially aggregated rodent counts once or twice per year: we provide here some support for the predation hypothesis from direct observations of mustelid and rodent individuals, at spatial and temporal scales commensurate to those of theoretical models. Surprisingly, the site level extinction probability of mustelids was found to increase when rodents were present. This is likely an artefact of the high extinction rate of rodents in the presence of mustelids. If a mustelid eradicates the rodents on a site within a primary occasion (i.e. within a week), and then leaves this site before the next primary occasion, this will appear as a mustelid extinction event conditional on the presence of rodents. This highlights a general challenge of multi-species occupancy models in terms of defining the length of primary occasions that are equally suitable for the different species included in the model and the interaction between them. This challenge is further highlighted by the lack of fit for the closed part of the model. Continuous time detection models, which may provide a solution to this issue, have recently been developed for single-season occupancy models (Kellner *et al*., 2022), but not yet been extended to dynamic state processes.

We note that numerous extensions to this model could be possible. Although we only included a simple binary covariate (season) for the detection probabilities in the case study, the model could be extended to include both temporal and spatial covariates on initial occupancy, colonization and extinction probabilities by including a multinomial logit link function following earlier multi-species occupancy models (Rota *et al*., 2016a; Fidino *et al*., 2019; Kéry & Royle, 2020). Our model also has the potential to be extended to more species (similar to models by Rota *et al*. 2016a and Fidino *et al*. 2019) or spatial levels. However, complicating the model further (e.g. by increasing number of species) may require regularization (shrinkage of some parameters) or variable selection (Hutchinson *et al*., 2015; McElreath, 2015).

To conclude, we developed a dynamic occupancy model for interacting species at two spatial scales, which estimates initial occupancy, colonization and extinction probabilities as well as detection probabilities. Applied to an Arctic rodent-mustelid system, the model provided evidence consistent with the predation hypothesis, while accounting for the fact that interactions between predators and prey arise from processes occurring at two nested spatial scales.

## 6 Acknowledgments

We thank H. Böhner for her invaluable help in classifying the camera trap images, M. Murphy, L.E. Støvern, I. Jensvoll and S. Killengreen for their help installing the camera trap monitoring design, M.L. Dahle and J.E. Knutsen for their help checking cameras and J.P. Mellard for language corrections. We thank the reviewers for constructive comments that considerably improved the manuscript. This work is a contribution from COAT (Climate-ecological Observatory for Arctic Tundra, www.coat.no), supported by the RCN (project nr. 245638), and COAT-Tools+, supported by Tromsø Research Foundation and UiT. OG was funded by the French National Research Agency (grant ANR-16-CE02-0007).

## 7 Author contributions

**Eivind F. Kleiven**: Conceptualization (equal); Data curation (supporting); Formal analysis (lead); Investigation (equal); Methodology (equal); Project administration (lead); Software (lead); Validation (equal); Visualization (lead); Writing - original draft (lead): Writing - review & editing (lead). **Frederic Barraquand**: Conceptualization (equal); Formal analysis (supporting); Investigation (equal); Methodology (equal); Project administration (supporting); Software (supporting); Validation (equal); Visualization (supporting); Writing - original draft (supporting); Writing - review & editing (equal). **Olivier Gimenez**: Conceptualization (equal); Formal analysis (supporting); Funding acquisition (lead); Methodology (supporting); Validation (supporting); Visualization (supporting); Writing - review & editing (equal). **John-André Henden**: Conceptualization (equal), Formal analysis (supporting); Methodology (supporting); Writing - review & editing (equal). **Rolf A. Ims**: Conceptualization (equal), Funding acquisition (lead); Investigation (equal); Writing - review & editing (equal). **Eeva M. Soininen**: Conceptualization (equal); Data Curation (lead); Funding acquisition (lead); Investigation (equal); Project administration (supporting); Visualization (supporting); Writing - original draft (supporting); Writing - review & editing (equal). **Nigel G. Yoccoz**: Conceptualization (equal), Funding acquisition (lead); Formal analysis (supporting); Methodology (supporting); Writing - review & editing (equal).

## 8 Data Availability

All data and code used in this manuscript are available at UiT Open Research Data (doi.org/10.18710/ZLW59W).

