## Appendix S1 for "A dynamic occupancy model for interacting species with two spatial scales"

Eivind F. Kleiven, Frédéric Barraquand, Olivier Gimenez, John-André Henden, Rolf A. Ims, Eeva M. Soininen & Nigel G. Yoccoz

#### 1 Initial occupancy state

It is necessary to define in what state the blocks and sites are in the first primary occasion (or initial state $\psi$). Since all observations are exclusively done at the site level and there is no information regarding the site state before the first sampling season, the initial latent site state ($z_{k,b,t=1}$) is modeled as the following categorical random variable (generalized Bernoulli), where $k$ denotes site, $b$ block and $t$ primary occasion

$$z_{k,b,t=1} \sim \text{Categorical}(\psi_b) \tag{1}$$

where $\psi_b$ is a vector describing the initial state probabilities of sites within a given block. To assure that the initial state probabilities sum to one, one state probability is obtained through subtraction. Then the initial state probability vector ($\psi_b$) can be written as

$$\psi_b = \begin{matrix} & \text{U} & \text{A} & \text{B} & \text{AB} \\ & [(1 - \psi_{A_b} - \psi_{B_b} - \psi_{AB_b}) & \psi_{A_b} & \psi_{B_b} & \psi_{AB_b}] \end{matrix} \tag{2}$$

where $\psi_{A_b}$, $\psi_{B_b}$ and $\psi_{AB_b}$ are the probabilities that a site within a given block will be in state A, B or AB respectively in the first primary occasion. Since there are no direct observations of the latent block state, it is assumed that the initial latent block state ($x_{b,t=1}$) is solely a function of the states of the sites within the given block (e.g. if at least one site is in state A and no sites are in state B or AB, then the block state is A).

We note that the initial state model presented above is rather simplistic. The reason for this is that the mustelid-rodent case study has a limited number of spatial replicates, compared to the relatively large number of primary occasions. Also the ecological interest in this example is mainly in the dynamical part of the model. In other cases where there is a stronger ecological interest in the initial states, we recommend implementing a initial state model where site state is dependent on block states, in line with the way we model transition probabilities.

#### 2 Simulation study

To investigate how detection and occupancy probability affect bias we simulate data from 6 sets of parameter values: 3 sets with varying detection probability and 3 sets with varying occupancy probability (see exact parameter values in Table S1). The parameters values for all scenarios were chosen to reflect a predator-prey setting, e.g. $\Gamma_A > \Gamma_{A|B}$ and $\epsilon_A < \epsilon_{A|B}$. Moreover, varying detection and occupancy scenarios was based on a review of occupancy studies by Specht *et al.* (2017) defining common species as occupancy $> 0.5$, rare species as occupancy $< 0.3$ and cryptic species as detection probability $< 0.3$.

| Parameter | $ld$ | $md$ | $hd$ | $lo$ | $mo$ | $ho$ |
|---|---|---|---|---|---|---|
| $\Gamma_A$ | 0.50 | 0.50 | 0.50 | 0.10 | 0.50 | 0.80 |
| $\Gamma_B$ | 0.10 | 0.10 | 0.10 | 0.05 | 0.10 | 0.20 |
| $\Gamma_{A\|B}$ | 0.05 | 0.05 | 0.05 | 0.05 | 0.05 | 0.20 |
| $\Gamma_{B\|A}$ | 0.40 | 0.40 | 0.40 | 0.20 | 0.40 | 0.70 |
| $E_A$ | 0.05 | 0.05 | 0.05 | 0.10 | 0.05 | 0.05 |
| $E_B$ | 0.60 | 0.60 | 0.60 | 0.60 | 0.60 | 0.80 |
| $E_{A\|B}$ | 0.50 | 0.50 | 0.50 | 0.50 | 0.50 | 0.40 |
| $E_{B\|A}$ | 0.20 | 0.20 | 0.20 | 0.20 | 0.20 | 0.20 |
| $p_A$ | 0.20 | 0.50 | 0.90 | 0.50 | 0.50 | 0.50 |
| $p_B$ | 0.10 | 0.50 | 0.80 | 0.50 | 0.50 | 0.50 |
| $\gamma_A$ | 0.50 | 0.50 | 0.50 | 0.30 | 0.50 | 0.80 |
| $\gamma_B$ | 0.30 | 0.30 | 0.30 | 0.30 | 0.30 | 0.30 |
| $\gamma_{A\|B}$ | 0.10 | 0.10 | 0.10 | 0.10 | 0.10 | 0.10 |
| $\gamma_{B\|A}$ | 0.70 | 0.70 | 0.70 | 0.60 | 0.70 | 0.70 |
| $\epsilon_A$ | 0.30 | 0.30 | 0.30 | 0.30 | 0.30 | 0.10 |
| $\epsilon_B$ | 0.80 | 0.80 | 0.80 | 0.80 | 0.80 | 0.60 |
| $\epsilon_{A\|B}$ | 0.90 | 0.90 | 0.90 | 0.90 | 0.90 | 0.60 |
| $\epsilon_{B\|A}$ | 0.10 | 0.10 | 0.10 | 0.10 | 0.10 | 0.10 |
| $\psi_{U,t=1}$ | 0.50 | 0.50 | 0.50 | 0.60 | 0.50 | 0.40 |
| $\psi_{A,t=1}$ | 0.25 | 0.25 | 0.25 | 0.20 | 0.25 | 0.30 |
| $\psi_{B,t=1}$ | 0.15 | 0.15 | 0.15 | 0.15 | 0.15 | 0.20 |
| $\psi_{AB,t=1}$ | 0.10 | 0.10 | 0.10 | 0.05 | 0.10 | 0.10 |

Table S1: Description of parameter values used in the simulation study under the 6 different scenarios. We use low, medium and high detection probability (ld, md, hd) and low, medium and high occupancy probability (lo, mo, ho).

We also plotted the trends in the simulated observed occupancy frequency at site level (Figure S1). This helps to clearly describe the actual data that was used, and show that the contrasted scenarios give the expected gradient in observed occupancy, where the low detection and low occupancy scenarios result in the lowest number of observations, while the medium detection and occupancy also gives an intermediate number of observation. High occupancy and detection result in the highest number of observations, so that the simulated datasets all serve their purpose.

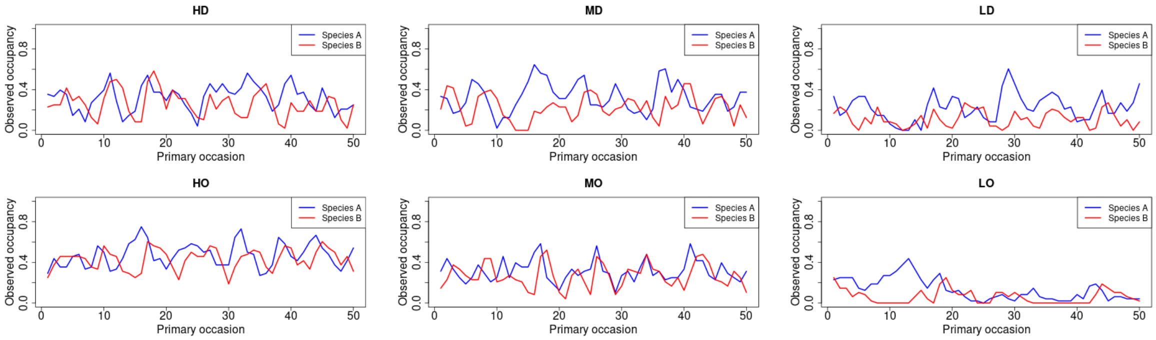


Figure S1: Description of the fraction of sites observed as occupied in different primary occasions (weeks) in the different simulation scenarios. The red line indicate the observed occupancy of species A, while the blue line indicate the observed occupancy of species B.

### 3 Camera trapping data

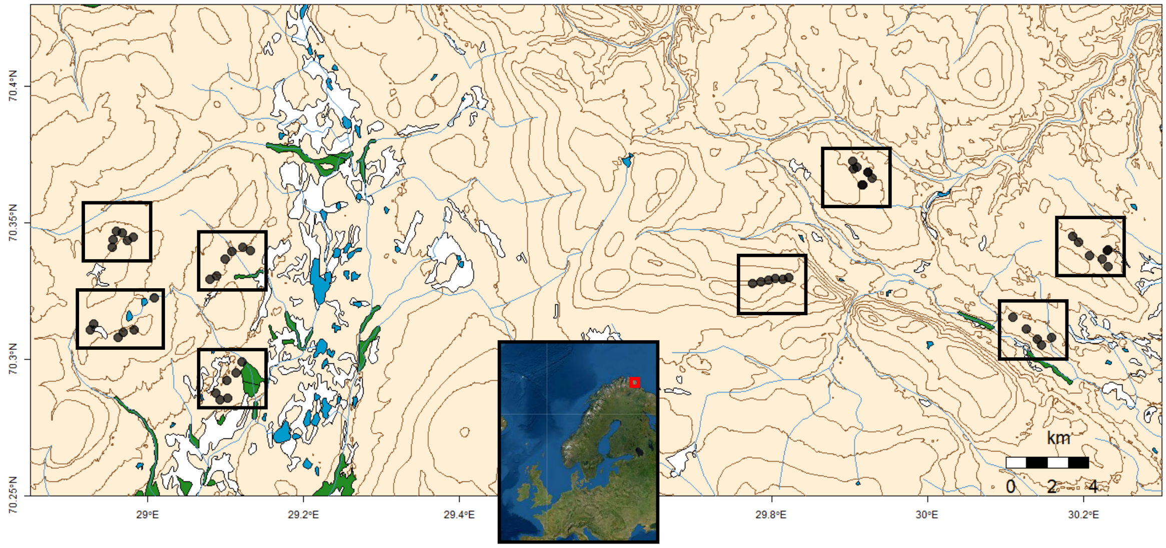


Figure S2: Map of the study area (Varanger peninsula, in northern Norway, red box), describing the sampling design of the empirical case study. Black dots mark camera trapping sites and black boxes highlight blocks of camera traps.

To illustrate the real world applicability of our model, we fit it to a camera trap dataset from the long-term monitoring program COAT (Climate-ecological Observatory of the Arctic Tundra) on small rodents (grey-sided vole, tundra vole and Norwegian lemming) and small mustelids (stoat and least weasel) on the Norwegian part of the low-arctic tundra. This monitoring program was started in autumn of 2015 and then consisted of 4 blocks of camera trap sites, with 11 sites with-in each block. In the summer of 2018 it was extended with 4 more blocks containing 12 sites within each block, to then make it a total of 8 blocks. The two functional groups of interest (i.e., small rodents and small mustelids) have different home ranges, so the spatial hierarchy in the sampling is designed so that sites (>300m apart) do not overlap the home range of the same individuals of small rodents, while they do so for small mustelids. The blocks are spaced out so that they do not overlap the home range of an individual of neither small rodents nor small mustelids (>3000m apart). At each site a camera trap consisting of a tunnel containing a motioned-triggered camera is placed in a natural runway for small mustelids and small rodents. The camera traps are out and active year-round and thus monitors the animals in continuous time (see Soininen *et al.* (2015) for further details about the camera traps). All camera trap images were automatically classified with the MLWIC R-package (Tabak *et al.*, 2019). The model was trained with a total of 47029 pictures from 8 different classes (see table S2 for details). The class "bad quality" was used for images where the quality is so low that it is likely that an animal could go undetected even though it was present (e.g. if the camera is full of snow or water). Such images were treated as missing observations in the data analysis. This training dataset was manually classified by experts (see Table S2 for classes and number of training pictures).

| Class ID | Number of pictures |
|---|---|
| Bad quality | 6695 |
| Bird | 1448 |
| Empty | 9001 |
| Least Weasel | 749 |
| Lemming | 8047 |
| Shrew | 8333 |
| Stoat | 3181 |
| vole | 9575 |
| Total | 47029 |

Table S2: Classes used for the training of the image classification model. In the right column, we show how many images were included in each category of the training data.

We used a separate test data consisting of 4425 expert classified images to validate the model. From these, the model classified 97.2% correctly.

The data were discretized into weeks (periods of 7 days) as primary occasions and days within a week as secondary occasions. We combine the data for the two functional groups into a multi-state occupancy data set with 4 states (U = none of the species are observer, A = only rodents observed, B = only mustelids observed or AB = both rodents and mustelids are observed). To analyze this data we included all weeks from the beginning of the monitoring in 2015 to 2021, making it a total of 304 weeks. Since more then half of the sites were only observed for the last 47 weeks the dataset includes much missing observations.

The present case study predator-prey system is strongly seasonal, with reproduction and population increase in summer, and population declines during the long snowy winter. In addition, there is inter-annual variation due to the typical 3 to 5-year population cycle in Arctic rodents (Ims & Fuglei, 2005). The seasonal and multi-annual components of dynamics are evident in the overall occupancy data (see Fig. S3).

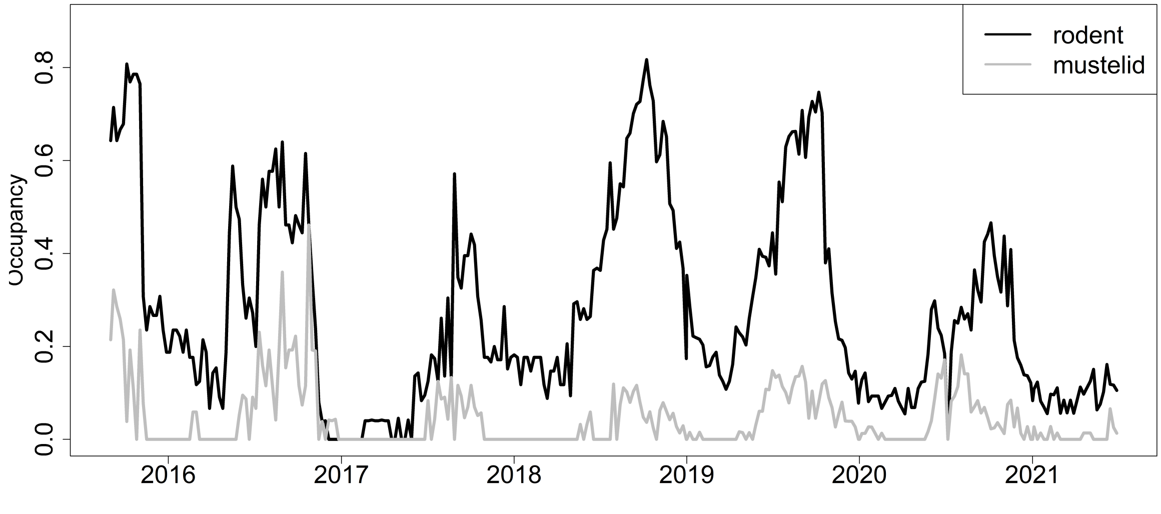


Figure S3: Proportion of active sites that were observed as occupied by either small rodents (black line) or small mustelids (gray line) in a given week (primary occasion). Note that the number of observed sites increased during the study as many sites were not observed in the first years
.

Table S3 gives the number of transitions observed between the block states (note that this is not the true number of transitions between latent states). It highlights that even though the case study covers many primary occasions, there are relatively few observed block level transitions, and especially so for states involving mustelids. We also see that some transitions are more common that others, something that potentially can give a hint about the data requirement of such a model.

| From site state | To site state: U | A | B | AB |
|---|---|---|---|---|
| $U$ | 699 | 94 | 26 | 1 |
| $A$ | 99 | 556 | 44 | 27 |
| $B$ | 26 | 43 | 107 | 22 |
| $AB$ | 1 | 27 | 19 | 20 |

Table S3: Number of observed block state transitions in the case study. State A denotes observed rodent presence and state B observed mustelid presence.

### 4 Model checking for case study

#### 4.1 Evaluating Goodness-of-fit

Explicit Goodness-of-fit (GOF) testing is challenging for occupancy models analysed in a Bayesian framework (Conn *et al.*, 2018). However, posterior predictive checks have been developed to investigate the GOF for simple dynamic occupancy models (Kéry & Royle, 2020) —other kinds of model checks, e.g. Dunn-Smyth residuals have been developed in a frequentist setup for static occupancy models (Warton *et al.*, 2017) or hidden Markov models. To assess the GOF in our Bayesian analyses, we adopted the approach presented by Kéry & Royle (2020). This approach considers separate fit statistics for the open (between primary occasions) and closed (within primary occasions) part of the model.

We tested the GOF on the site level and did separate tests for the two species (i.e. functional groups). To do this we considered the occupancy model for interacting species with two spatial scales to provide information on the latent occupancy state of the two species separately. That is, we considered the latent state of species A ($z_A$) to be 1 if the model estimated the latent state of a site ($z$) as A or AB, and similarly for latent state of species B ($z_B$) if latent state of a site was estimated to $B$ or $AB$. We perform the GOF-test as a posterior predictive check, where we simulate data under the fitted model and compare it with the observed data. For the test for the open part of the model we compared the number of observed transitions (i.e. the number of site level state transitions in the observed dataset $y_{b,k,t,j}$ as well as the dataset simulated under the fitted model) to the number of expected transitions as estimated from the model parameters (e.g. the expected number of times a site will remain unoccupied by species A is calculated as $\sum_{k=1}^{BK}(1-z_{A_{k,t}})\left((1-z_{B_{k,t}})(1-\hat{\gamma}_A)+z_{B_{k,t}}(1-\hat{\gamma}_{AB})\right)$ for each primary occasion).

For the closed part of the model, observed detection frequencies (i.e. frequencies of observations within a primary occasion) and expected detection frequencies calculated from the estimated model parameters (i.e. $z_{A_{k,t}}p_t$ for each secondary occasions $j$) at individual sites were compared. To compare observed and expected values for both the real dataset and the dataset simulated under the fitted model we calculated $\chi^2$ discrepancies as follows:

$$\chi^2_{\text{obs}} = \sum_{t=1}^{T-1}\sum_{i,i'\in\{1,2\}}(O^{\text{obs}}_{i,i',t}-E_{i,i',t})^2/E_{i,i',t} \tag{3}$$

$$\chi^2_{\text{rep}} = \sum_{t=1}^{T-1}\sum_{i,i'\in\{1,2\}}(O^{\text{rep}}_{i,i',t}-E_{i,i',t})^2/E_{i,i',t} \tag{4}$$

where $O^{\text{obs}}_{t,i,i'}$ is the number of transitions between two consecutive primary occasions from state $i$ to $i'$. Transitions occur between observed not latent states (observed states being defined in the dataset by $y_{b,k,t,j}$), so that $O^{\text{rep}}_{i,i',t}$ is the number of transitions between observed states of two consecutive primary occasions in a replicated dataset which is simulated under the fitted model. The states $(z_A, z_B)$ can take on two values for each species, $U$ or $A$ vs $U$ or $B$, which is coded in the $O$ and $E$ matrices as 1 and 2. $E_{t,i,i'}$ is the number of transitions between states expected from the model. We compute the same quantities for frequency (number of times a species is seen within a primary occasion) in place of transitions. We note that while $O^{\text{obs}}_{i,i',t}$ and $O^{\text{rep}}_{i,i',t}$ are calculated exactly, $E_{i,i',t}$ are approximated assuming that all sites within a block might be colonized at any time point independent of the actual block state.

For the posterior predictive check, discrepancies from the observed dataset are compared to discrepancies in the dataset simulated under the model. Bayesian $p$-values were calculated as the fraction of the MCMC-samples for which the discrepancies for the dataset simulated under the fitted model were larger than the discrepancy in the real dataset (i.e. the proportion of points above the 1:1 line in a plot of the $\chi^2$ discrepancies for the expected and the observed data as in Fig. S5). Bayesian $p$-values close to 0.5 imply a good fit. However, there is still some debate about exactly how close to 0.5 the Bayesian $p$-value needs to be to assume a good fit (Gelman, 2013). Therefore, we first applied the GOF-test on a simulated data set (mid occupancy scenario from the simulation study) to investigate how the test would preform under ideal circumstances (see Fig. S4). However, we do stress that this is not a complete investigation of how this GOF-test performs for complex occupancy models, but rather a simple verification of what the posterior predictive check might look like under ideal circumstances. Indeed, we want to emphasize that more work is needed to understand the sensitivity of GOF-tests for complex occupancy models and how the outputs of these tests should be interpreted (Kéry & Royle, 2020).

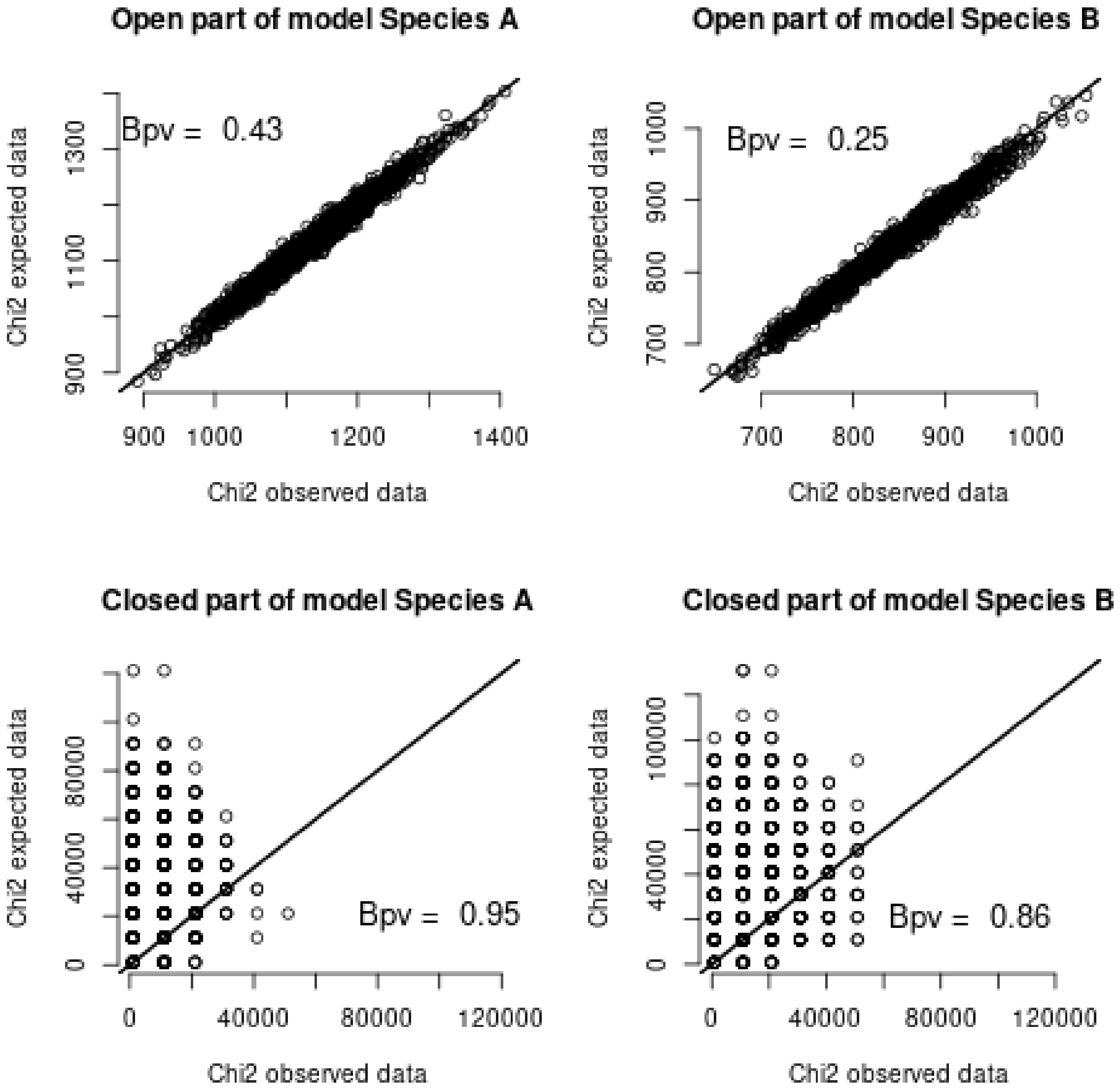


Figure S4: Posterior predictive check of the open and closed parts of the model based on data from the simulation study (medium detection scenario) for the two different species groups. Each point represent one sample from the posterior distribution. The Bayesian p-value (Bpv) is the proportion of points above the 1:1 line.

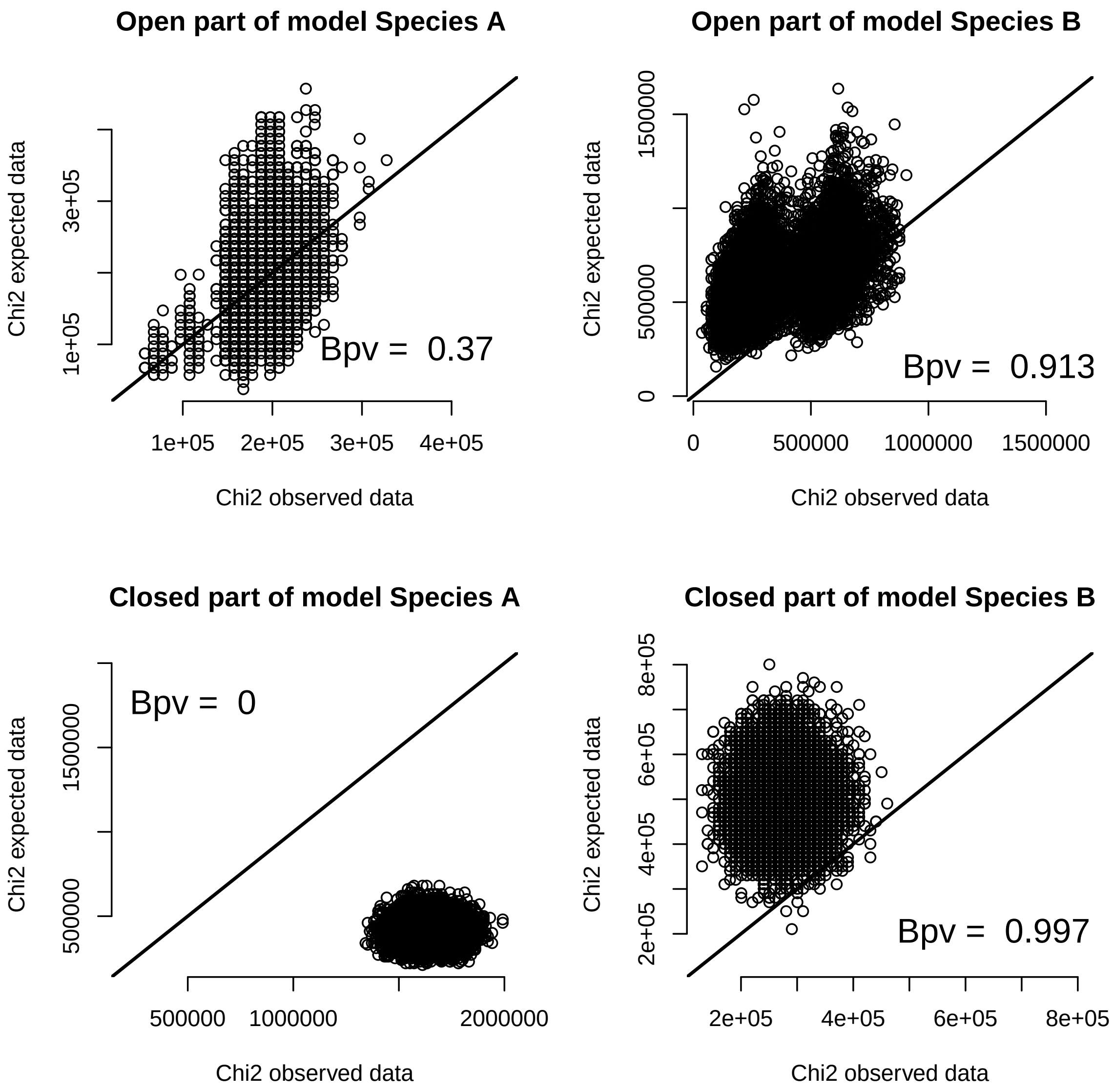


Figure S5: Posterior predictive check of the open and closed parts of the model based on the case study data for the two different species groups (rodents (A) and mustelids (B)). Each point represent one sample from the posterior distribution. The Bayesian p-value (Bpv) is the proportion of points above the 1:1 line.

We see from figure S4 and S5 that regarding the case study data, the $\chi^2$ discrepancies between the replicated and observed data are a little more different from each other than for data simulated under the theoretical model. There is a large discrepancy for species A, and to a lesser extent for species B, in the closed part of the model. This could indicate that there are factors affecting the detection probabilities that we do not account for. However, regarding the open part of the model for both functional groups we see that there is not a considerable difference between what is expected under ideal circumstances (data simulated under the model) and what we see with the case study data.

#### 4.2 Prior sensitivity analysis

To ensure that the parameters were not too dependent on the prior distribution, we performed a prior sensitivity analysis. This was done by running the model with three different sets of priors (see table S4) and comparing parameter estimates. This prior sensitivity analysis can also be used to investigate parameter identifiability by estimating the prior-posterior overlap.

| Parameter | previously used priors | centered prior set | skewed prior set |
|---|---|---|---|
| $\Gamma_A$ | Uniform(0,1) | Beta(4,4) | Beta(2,4) |
| $\Gamma_{A\|B}$ | Uniform(0,1) | Beta(4,4) | Beta(2,4) |
| $\Gamma_B$ | Uniform(0,1) | Beta(4,4) | Beta(2,4) |
| $\Gamma_{B\|A}$ | Uniform(0,1) | Beta(4,4) | Beta(2,4) |
| $E_A$ | Uniform(0,1) | Beta(4,4) | Beta(2,4) |
| $E_{A\|B}$ | Uniform(0,1) | Beta(4,4) | Beta(4,2) |
| $E_B$ | Uniform(0,1) | Beta(4,4) | Beta(4,2) |
| $E_{B\|A}$ | Uniform(0,1) | Beta(4,4) | Beta(2,4) |
| $\gamma_A$ | Uniform(0,1) | Beta(4,4) | Beta(4,2) |
| $\gamma_{A\|B}$ | Uniform(0,1) | Beta(4,4) | Beta(2,4) |
| $\gamma_B$ | Uniform(0,1) | Beta(4,4) | Beta(2,4) |
| $\gamma_{B\|A}$ | Uniform(0,1) | Beta(4,4) | Beta(4,2) |
| $\epsilon_A$ | Uniform(0,1) | Beta(4,4) | Beta(2,4) |
| $\epsilon_{A\|B}$ | Uniform(0,1) | Beta(4,4) | Beta(4,2) |
| $\epsilon_B$ | Uniform(0,1) | Beta(4,4) | Beta(4,2) |
| $\epsilon_{B\|A}$ | Uniform(0,1) | Beta(4,4) | Beta(2,4) |
| $\alpha_{A0}$ | Normal(0,1) | logistic(0,1) | Normal(0.5,1) |
| $\alpha_{B0}$ | Normal(0,1) | logistic(0,1) | Normal(-0.5,1) |
| $\psi_1$ | Uniform(0,0.5) | Beta(4,4) | Beta(2,4) |
| $\psi_2$ | Uniform(0,0.5) | Beta(4,4) | Beta(2,4) |
| $\psi_2$ | Uniform(0,0.5) | Beta(4,4) | Beta(2,4) |

Table S4: Priors used for the prior sensitivity analysis. $\alpha_{A0}$ and $\alpha_{B0}$ are the intercept in the logit link function for the detection probabilities $p_A$ and $p_B$. These parameters are on the logit scale. $\psi_1$, $\psi_2$ and $\psi_3$ are the initial state probabilities for states A, B and AB. $\Gamma$ and $E$ are the colonization and extinction probabilities on the block level while $\gamma$ and $\epsilon$ are the colonization and extinction probabilities on the site level. All initial state, colonization and extinction probabilities are naturally bounded between zero and one. Exact definitions of these with subscripts can be found in Tables 1 and 2 in the main text.

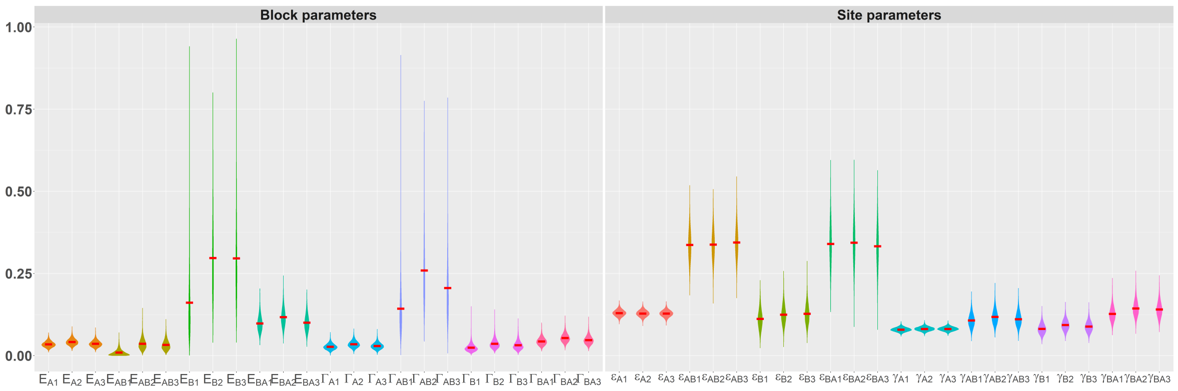


Figure S6: Violin plot of colonization and extinction probability estimates with the different sets of priors. The red bar indicates the posterior mean. In the x-axis labels 1-3 indicate the prior set used.

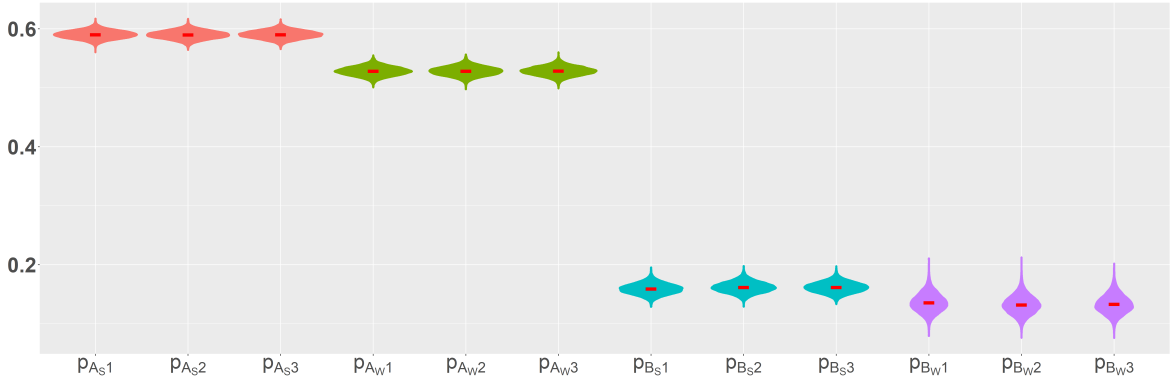


Figure S7: Violin plot of the detection probabilities estimates with the different sets of priors. The red bar indicates the posterior mean. In the x-axis labels 1-3 indicate the prior set used.

We see from figure S6 and S7 that the estimates of detection, colonization and extinction probabilities seem to have little sensitivity to the prior distributions, except for $\Gamma_{A|B}$ and $E_B$ where the posterior seems to change slightly depending on the prior distribution.

#### 4.3 Parameter identifiability

To check parameter identifiability we investigated the overlap between prior and posterior distributions. This was done for all colonization, extinction and detection parameters for the same 3 sets of prior distributions that were used in the prior sensitivity analysis (see table S4). From Figures S8, S9 and S10 we see that the estimated posterior distributions for all parameters are estimated similarly for the 3 sets of priors and that they are consistently different from the prior distributions. Therefore, it appears that all colonization and extinction parameters on both spatial level are identifiable from the data.

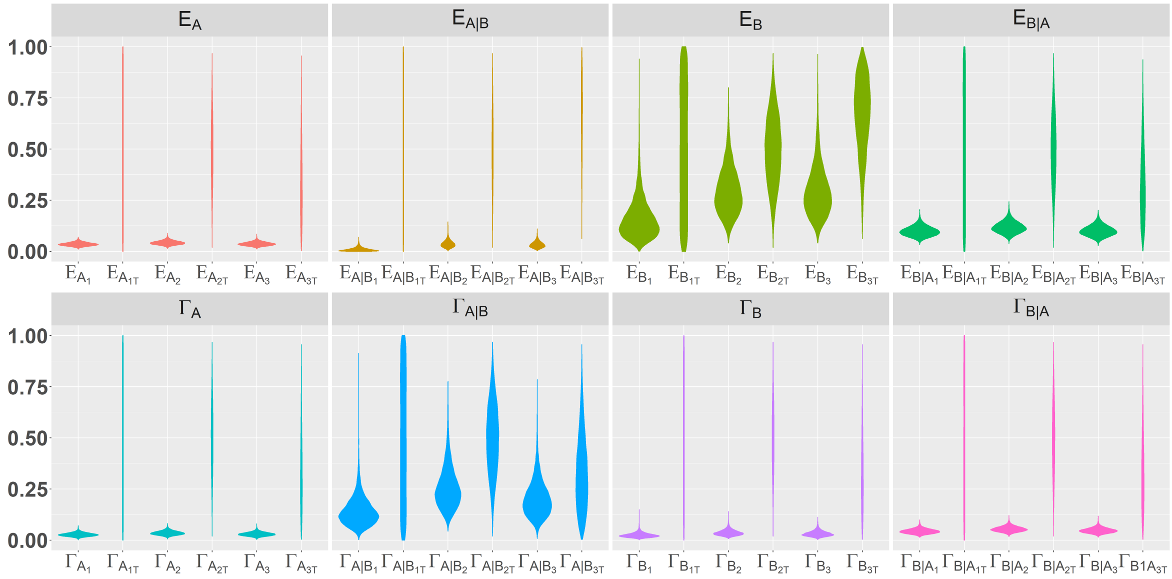


Figure S8: Violin plot of both prior (indicated by subscript T) and posterior distributions for block colonization and extinction parameters with the three sets of priors. Subscripts 1 to 3 indicate the prior set used.

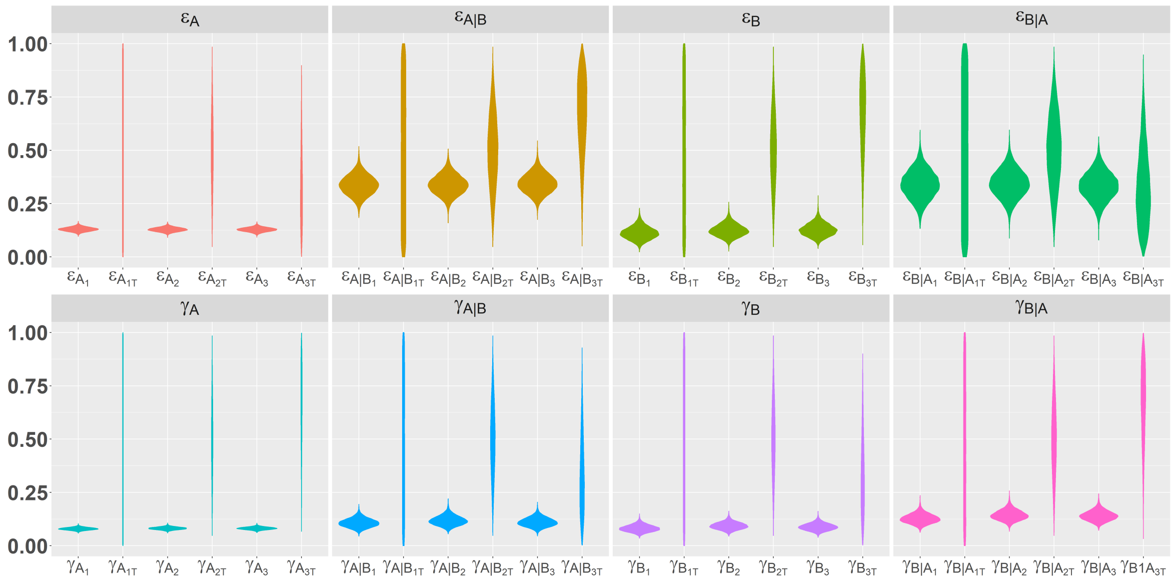


Figure S9: Violin plot of both prior (indicated by subscript T) and posterior distributions for site colonization and extinction parameters with the three sets of priors. Subscripts 1 to 3 indicate the prior set used.

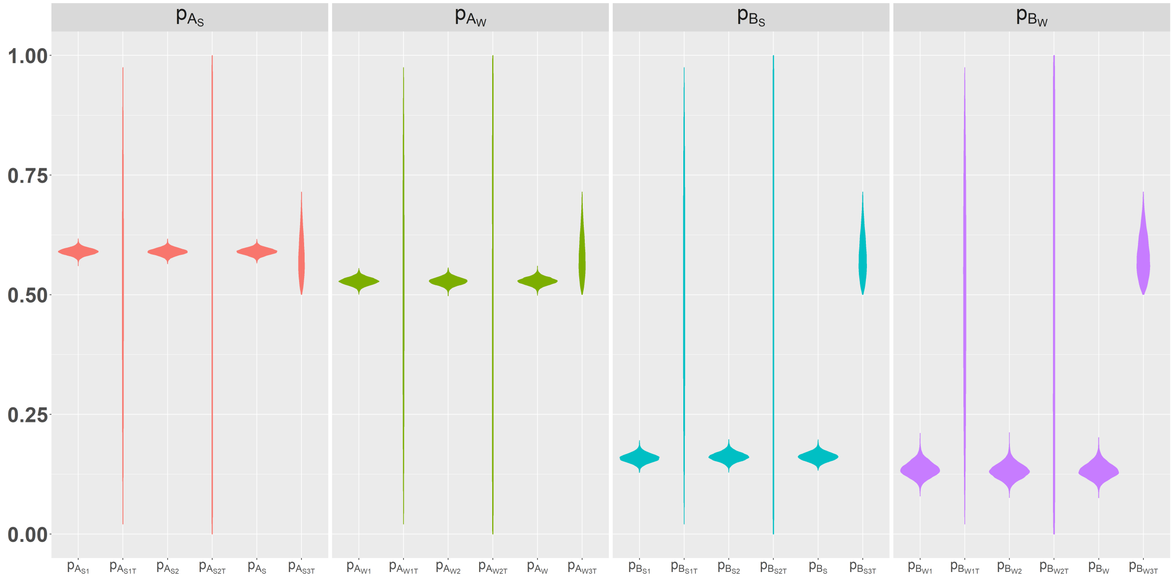


Figure S10: Violin plot of both prior (indicated by subscript T) and posterior distributions for block colonization and extinction parameters with the three sets of priors. Subscript S and W refer to summer and winter estimates, while 1 to 3 indicate the prior set used.

### 5 Results

#### 5.1 Simulation study

In this section we give a more detailed description of the simulation results.

Tables S7 and S6 give the detailed results from the simulation study. We see that the model is able to estimate most parameters without bias. Some small biases do appear at times, but mostly under the low detection probability scenario.

| | low det | | | mid det | | | high det | | |
|---|---|---|---|---|---|---|---|---|---|
| Parameter | true | mean | sd | true | mean | sd | true | mean | sd |
| $\Gamma_A$ | 0.50 | 0.43 | 0.06 | 0.50 | 0.48 | 0.04 | 0.50 | 0.49 | 0.05 |
| $\Gamma_{A\|B}$ | 0.05 | 0.05 | 0.03 | 0.05 | 0.07 | 0.03 | 0.05 | 0.06 | 0.02 |
| $\Gamma_B$ | 0.10 | 0.06 | 0.03 | 0.1 | 0.10 | 0.03 | 0.10 | 0.10 | 0.03 |
| $\Gamma_{B\|A}$ | 0.40 | 0.36 | 0.04 | 0.4 | 0.40 | 0.05 | 0.40 | 0.41 | 0.04 |
| $E_A$ | 0.05 | 0.10 | 0.03 | 0.05 | 0.06 | 0.02 | 0.05 | 0.06 | 0.02 |
| $E_{A\|B}$ | 0.50 | 0.61 | 0.08 | 0.50 | 0.53 | 0.05 | 0.50 | 0.52 | 0.06 |
| $E_B$ | 0.60 | 0.73 | 0.08 | 0.60 | 0.63 | 0.06 | 0.60 | 0.60 | 0.05 |
| $E_{B\|A}$ | 0.20 | 0.21 | 0.04 | 0.20 | 0.20 | 0.04 | 0.20 | 0.20 | 0.04 |
| $\gamma_A$ | 0.50 | 0.46 | 0.02 | 0.50 | 0.49 | 0.02 | 0.50 | 0.50 | 0.01 |
| $\gamma_{A\|B}$ | 0.10 | 0.06 | 0.03 | 0.10 | 0.08 | 0.03 | 0.10 | 0.09 | 0.03 |
| $\gamma_B$ | 0.30 | 0.23 | 0.05 | 0.30 | 0.29 | 0.02 | 0.30 | 0.30 | 0.02 |
| $\gamma_{B\|A}$ | 0.70 | 0.74 | 0.07 | 0.70 | 0.70 | 0.02 | 0.70 | 0.70 | 0.02 |
| $\epsilon_A$ | 0.30 | 0.34 | 0.03 | 0.30 | 0.31 | 0.02 | 0.30 | 0.30 | 0.01 |
| $\epsilon_{A\|B}$ | 0.90 | 0.94 | 0.03 | 0.90 | 0.92 | 0.03 | 0.90 | 0.91 | 0.03 |
| $\epsilon_B$ | 0.80 | 0.84 | 0.07 | 0.80 | 0.81 | 0.02 | 0.80 | 0.80 | 0.02 |
| $\epsilon_{B\|A}$ | 0.10 | 0.11 | 0.04 | 0.10 | 0.09 | 0.02 | 0.10 | 0.09 | 0.02 |

Table S5: Results from the 3 scenarios with varying detection probabilities in the simulation study given as mean and standard deviation of the posterior means from the 50 replicated sets.

| | low occ | | | mid occ | | | high occ | | |
|---|---|---|---|---|---|---|---|---|---|
| Parameter | true | mean | sd | true | mean | sd | true | mean | sd |
| $\Gamma_A$ | 0.10 | 0.10 | 0.02 | 0.50 | 0.50 | 0.05 | 0.80 | 0.79 | 0.05 |
| $\Gamma_{A\|B}$ | 0.05 | 0.07 | 0.03 | 0.05 | 0.06 | 0.03 | 0.20 | 0.21 | 0.05 |
| $\Gamma_B$ | 0.05 | 0.05 | 0.01 | 0.10 | 0.10 | 0.03 | 0.20 | 0.19 | 0.05 |
| $\Gamma_{B\|A}$ | 0.20 | 0.22 | 0.05 | 0.40 | 0.40 | 0.06 | 0.70 | 0.69 | 0.04 |
| $E_A$ | 0.10 | 0.10 | 0.03 | 0.05 | 0.06 | 0.02 | 0.05 | 0.07 | 0.02 |
| $E_{A\|B}$ | 0.50 | 0.53 | 0.07 | 0.50 | 0.53 | 0.05 | 0.40 | 0.39 | 0.04 |
| $E_B$ | 0.60 | 0.63 | 0.08 | 0.60 | 0.62 | 0.06 | 0.80 | 0.79 | 0.05 |
| $E_{B\|A}$ | 0.20 | 0.21 | 0.06 | 0.20 | 0.21 | 0.04 | 0.20 | 0.19 | 0.03 |
| $\gamma_A$ | 0.30 | 0.29 | 0.02 | 0.50 | 0.49 | 0.02 | 0.80 | 0.79 | 0.01 |
| $\gamma_{A\|B}$ | 0.10 | 0.10 | 0.04 | 0.10 | 0.09 | 0.03 | 0.10 | 0.08 | 0.02 |
| $\gamma_B$ | 0.30 | 0.29 | 0.02 | 0.30 | 0.29 | 0.02 | 0.30 | 0.29 | 0.02 |
| $\gamma_{B\|A}$ | 0.60 | 0.59 | 0.04 | 0.70 | 0.70 | 0.02 | 0.70 | 0.69 | 0.02 |
| $\epsilon_A$ | 0.30 | 0.30 | 0.03 | 0.30 | 0.31 | 0.02 | 0.10 | 0.10 | 0.01 |
| $\epsilon_{A\|B}$ | 0.90 | 0.90 | 0.05 | 0.90 | 0.92 | 0.02 | 0.60 | 0.63 | 0.03 |
| $\epsilon_B$ | 0.80 | 0.80 | 0.04 | 0.80 | 0.80 | 0.02 | 0.60 | 0.61 | 0.03 |
| $\epsilon_{B\|A}$ | 0.10 | 0.10 | 0.04 | 0.10 | 0.09 | 0.02 | 0.10 | 0.09 | 0.02 |

Table S6: Results from the 3 scenarios with varying occupancy probabilities in the simulation study given as mean and standard deviation of the posterior means from the 50 replicated sets.

| | low det | | mid det | | high det | | low occ | | mid occ | | high occ | |
|---|---|---|---|---|---|---|---|---|---|---|---|---|
| Par | bias | rel. | bias | rel. | bias | rel. | bias | rel. | bias | rel. | bias | rel. |
| $\Gamma_A$ | 0.07 | 14% | 0.02 | 4% | 0.01 | 2% | 0 | 0 | 0 | 0 | 0.01 | 1% |
| $\Gamma_{A\|B}$ | 0 | 0 | 0.02 | 40% | 0.01 | 20% | 0.02 | 40% | 0.01 | 20% | 0.01 | 5% |
| $\Gamma_B$ | 0.04 | 40% | 0 | 0% | 0 | 0% | 0 | 0% | 0 | 0% | 0.01 | 5% |
| $\Gamma_{B\|A}$ | 0.04 | 10% | 0 | 0% | 0.01 | 3% | 0.02 | 10% | 0 | 0% | 0.01 | 1% |
| $E_A$ | 0.05 | 50% | 0.01 | 20% | 0.01 | 20% | 0 | 0% | 0.01 | 20% | 0.02 | 40% |
| $E_{A\|B}$ | 0.11 | 22% | 0.03 | 6% | 0.02 | 4% | 0.03 | 6% | 0.03 | 6% | 0.01 | 3% |
| $E_B$ | 0.13 | 22% | 0.03 | 5% | 0 | 0% | 0.03 | 5% | 0.02 | 3% | 0.01 | 1% |
| $E_{B\|A}$ | 0.01 | 5% | 0 | 0% | 0 | 0% | 0.01 | 5% | 0.01 | 5% | 0.01 | 5% |
| $\gamma_A$ | 0.04 | 8% | 0.02 | 4% | 0 | 0% | 0.01 | 3% | 0.01 | 2% | 0.01 | 1% |
| $\gamma_{A\|B}$ | 0.04 | 40% | 0.02 | 20% | 0.01 | 10% | 0 | 0% | 0.01 | 10% | 0.02 | 20% |
| $\gamma_B$ | 0.07 | 23% | 0.01 | 3% | 0 | 0% | 0.01 | 3% | 0.01 | 3% | 0.01 | 3% |
| $\gamma_{B\|A}$ | 0.04 | 6% | 0 | 0% | 0 | 0% | 0.01 | 1% | 0 | 0% | 0.01 | 1% |
| $\epsilon_A$ | 0.04 | 13% | 0.01 | 3% | 0 | 0% | 0 | 0% | 0.01 | 3% | 0 | 0% |
| $\epsilon_{A\|B}$ | 0.04 | 4% | 0.02 | 2% | 0.01 | 1% | 0 | 0% | 0.02 | 2% | 0.03 | 5% |
| $\epsilon_B$ | 0.04 | 5% | 0.01 | 1% | 0 | 0% | 0 | 0% | 0 | 0% | 0.01 | 2% |
| $\epsilon_{B\|A}$ | 0.01 | 10% | 0.01 | 10% | 0.01 | 10% | 0 | 0% | 0.01 | 10% | 0.01 | 10% |

Table S7: Comparison of the true and estimated parameter values in the simulation study. The table gives the absolute and relative bias.

From Figure S11 and S12 we see that detection probabilities ($p_A$ and $p_B$) and initial occupancy probabilities ($\psi_{A_{t-1}}$, $\psi_{B_{t-1}}$ and $\psi_{AB_{t-1}}$) are estimated without bias.

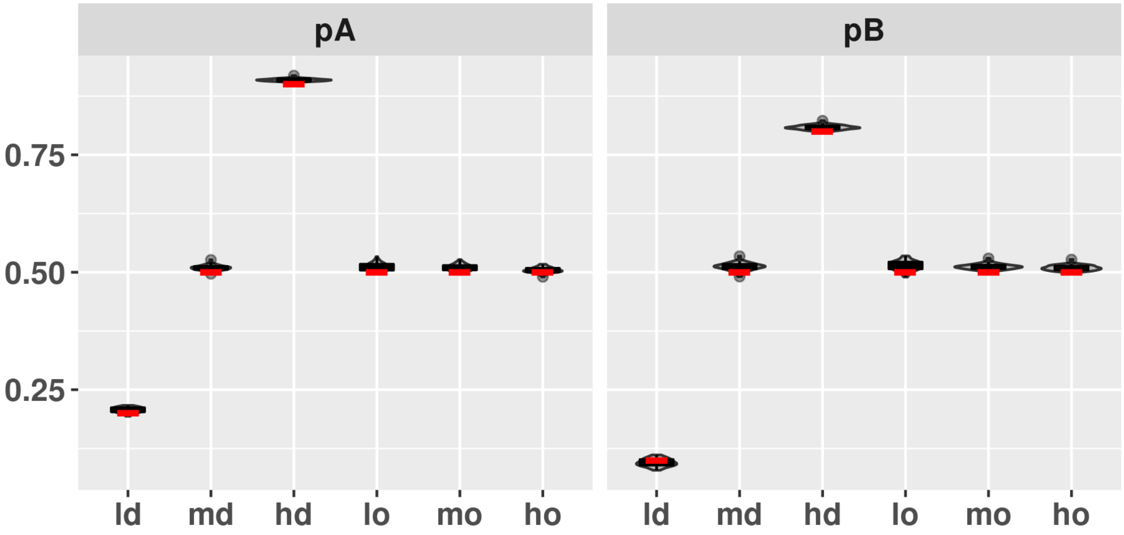


Figure S11: Violin plot and boxplot of the posterior mean of detection probability ($p_A$ and $p_B$) from the 50 simulations. The red bar indicates the true parameter values. x-axis displays the 6 different data scenarios (low, medium and high detection probability: ld, md, hd, and low, medium and high occupancy probability: lo, mo,ho).

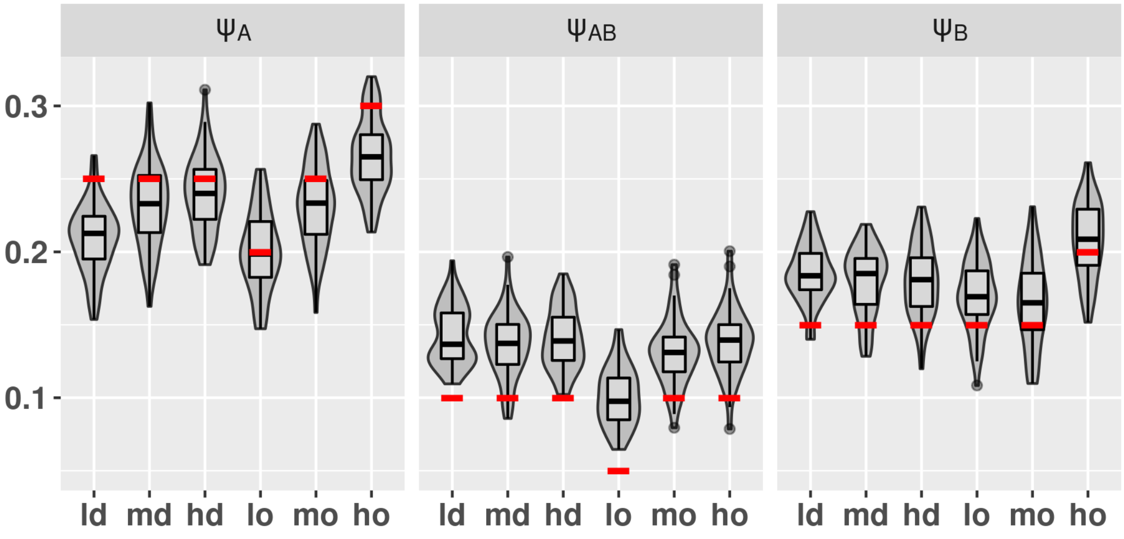


Figure S12: Violin plot and boxplot of the posterior mean of initial occupancy ($\psi_{t=1}$) from the 50 simulations. The red bar indicates the true parameter values. x-axis displays the 6 different data scenarios (low, medium and high detection probability: ld, md, hd, and low, medium and high occupancy probability: lo, mo,ho).

#### 5.2 Case study

This section gives detailed results on the case study.

Table S8 gives more details on the estimated colonization and extinction parameters, while table S9 gives more information on the estimated differences between colonization and extinction parameters depending on the presence or absence of the other species. Figure S13 displays the difference in detection probability between the two species groups and the two seasons as described in the main text.

| Parameter | mean | sd | 95% CI |
|---|---|---|---|
| $\Gamma_A$ | 0.027 | 0.009 | 0.012 0.046 |
| $\Gamma_{A\|B}$ | 0.143 | 0.072 | 0.045 0.313 |
| $\Gamma_B$ | 0.025 | 0.011 | 0.009 0.052 |
| $\Gamma_{B\|A}$ | 0.044 | 0.012 | 0.024 0.069 |
| $E_A$ | 0.035 | 0.009 | 0.020 0.054 |
| $E_{A\|B}$ | 0.010 | 0.009 | 0.001 0.034 |
| $E_B$ | 0.161 | 0.098 | 0.036 0.410 |
| $E_{B\|A}$ | 0.098 | 0.024 | 0.057 0.149 |
| $\gamma_A$ | 0.079 | 0.006 | 0.068 0.092 |
| $\gamma_{A\|B}$ | 0.108 | 0.021 | 0.071 0.151 |
| $\gamma_B$ | 0.082 | 0.016 | 0.054 0.115 |
| $\gamma_{B\|A}$ | 0.128 | 0.022 | 0.088 0.174 |
| $\epsilon_A$ | 0.130 | 0.010 | 0.112 0.149 |
| $\epsilon_{A\|B}$ | 0.337 | 0.046 | 0.249 0.428 |
| $\epsilon_B$ | 0.112 | 0.029 | 0.060 0.175 |
| $\epsilon_{B\|A}$ | 0.340 | 0.066 | 0.212 0.472 |

Table S8: Results from the case study given as mean and standard deviation from the posterior distribution in addition to the 95% credible intervals.

| Parameter | mean | 95% CI |
|---|---|---|
| $\gamma_A$ - $\gamma_{A\|B}$ | -0.027 | -0.065 0.006 |
| $\gamma_B$ - $\gamma_{B\|A}$ | -0.045 | -0.089 -0.005 |
| $\epsilon_A$ - $\epsilon_{A\|B}$ | - 0.207 | -0.288 -0.129 |
| $\epsilon_B$ - $\epsilon_{B\|A}$ | -0.228 | -0.347 -0.111 |
| $\Gamma_A$ - $\Gamma_{A\|B}$ | -0.104 | -0.247 -0.024 |
| $\Gamma_B$ - $\Gamma_{B\|A}$ | -0.019 | -0.045 0.008 |
| $E_A$ - $E_{A\|B}$ | 0.026 | 0.002 0.044 |
| $E_B$ - $E_{B\|A}$ | 0.043 | -0.063 0.257 |

Table S9: Estimated differences between dependent and independent colonization and extinction probabilities with 95% credible intervals.

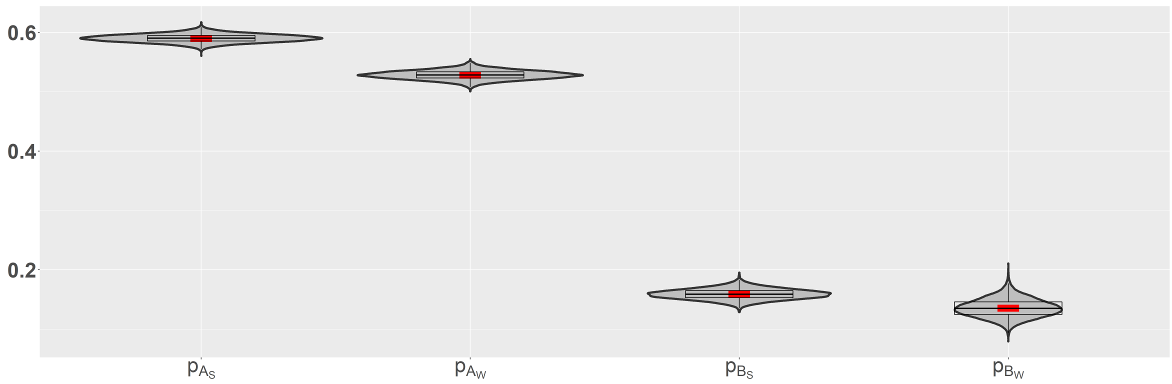


Figure S13: Violin plot of the posterior distribution of detection probability ($p_A$ and $p_B$) for summer (S) and winter (W) in black. The red bar indicates the mean of the posterior distribution.

Figure S14 shows the estimated initial occupancy probabilities from the case study ($\psi_{t=1}$). We see some variation between blocks, where blocks b2 and b4 have a higher occupancy. Note that $\psi_1$ is occupancy probability of species A only, $\psi_2$ is occupancy probability of species B only and $\psi_3$ is the probability that both species A and B occupy a given site. Block b2 has high occupancy probability of species A ($\psi_1$) and b4 has high occupancy probability of species B ($\psi_3$). Please also note that we do not have any observations from blocks b6-b8 in the initial season, and that we do not include any spatial covariates in the model.

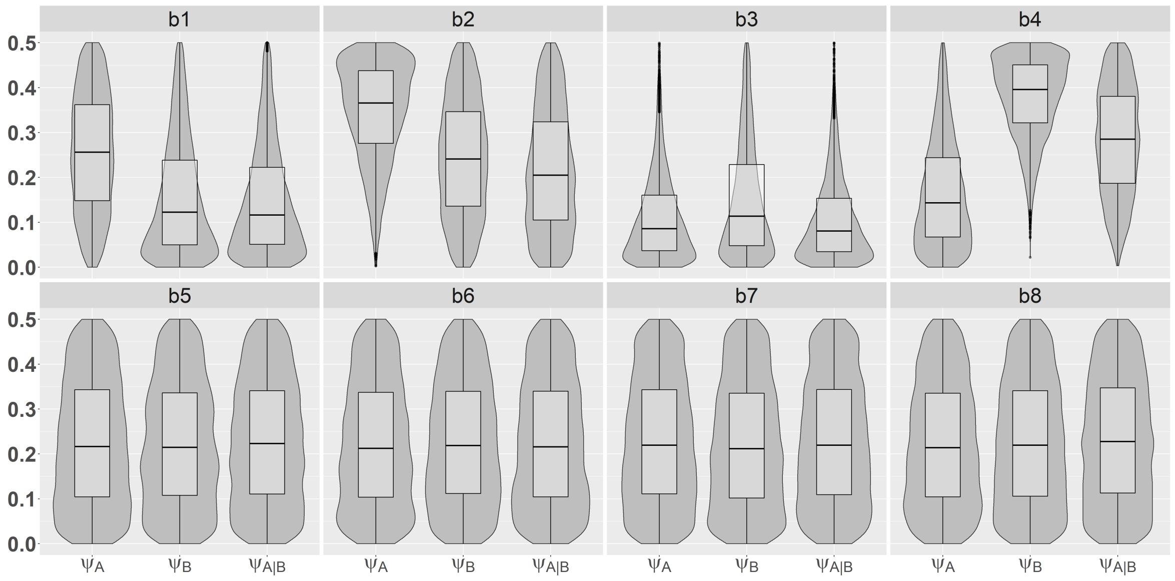


Figure S14: Violin plot and boxplot of the posterior mean of initial occupancy ($\psi_{t=1}$) for each of the 8 blocks (b1-b8).
